# *In vivo* cellular localisation and nanoscale organisation of NLRP3 inflammasomes

**DOI:** 10.64898/2026.08.06.743215

**Authors:** Christopher Hoyle, Benjamin Llewellyn, Helen Parker, Katie Murray, Andrew D. Greenhalgh, Jonathan D. Worboys, Rodrigo Diaz Pino, Julia Ogden, Adam Johnson, Antony D. Adamson, Kevin N. Couper, Catherine B Lawrence, Gloria Lopez-Castejon, Martin Lowe, David Brough, Jack P. Green

## Abstract

The NLRP3 inflammasome is a critical regulator of inflammation, yet the localisation, organisation, and cellular sources of endogenous NLRP3 inflammasomes remain incompletely understood. Here, we generated NLRP3-mScarlet-I endogenous reporter mice enabling visualisation of NLRP3 at physiological levels in primary cells and *in vivo*. We show that activated NLRP3 associated with PI4P-positive membranes from multiple organelles, supporting a model where diverse membrane platforms act as a scaffold to nucleate inflammasome assembly. Super-resolution imaging revealed that NLRP3 and ASC occupy distinct nanoscale architectures within the inflammasome, with NLRP3 displaying marked structural heterogeneity and stimulus-dependent organisation. Unexpectedly, circulating monocytes and neutrophils, rather than tissue-resident populations, emerged as the dominant NLRP3-expressing cells *in vivo* which rapidly infiltrated tissues following systemic inflammation, highlighting an underappreciated cellular source of rapid inflammasome-driven responses. These findings reveal previously unrecognised insights into inflammasome organisation and localisation, establishing a powerful resource for investigating endogenous NLRP3 biology in health and disease.

## Main

NLR family pyrin domain containing 3 (NLRP3) is a cytosolic protein expressed by immune cells and is important for host responses to infection and injury. Tight regulation of NLRP3 activation is essential for maintaining physiological responses to infection and injury; however, aberrant or excessive NLRP3 activity underpins the development of numerous inflammatory, cardiovascular and neurodegenerative diseases (1). A central feature of NLRP3 is its ability to sense alterations in cellular homeostasis induced by pathogen- and damage-associated molecular patterns (PAMPs and DAMPs, respectively) to trigger the assembly of inflammasome complexes that catalyse the activation and secretion of interleukin (IL)-1 family cytokines, namely IL-1β and IL-18 (2). Once activated, NLRP3 interacts with the adaptor protein apoptosis-associated speck-like protein containing a CARD (ASC) that subsequently recruits and activates the protease caspase-1 that cleaves pro forms of IL-1β and IL-18 into their active secreted forms. Caspase-1 also cleaves gasdermin D, which then forms membrane pores, serving as the conduit for IL-1 secretion, and drives a ninjurin-1-dependent inflammatory mode of cell death called pyroptosis (3). NLRP3-driven inflammation is implicated in many major diseases and there is great interest in understanding its biology to develop new treatments (4).

There are several current theories for how NLRP3 is activated. NLRP3 is reported to be recruited to vesicles of dispersed *trans*-Golgi network (dTGN) through an interaction between its polybasic motif and the lipid phosphatitdylinositol-4-phosphate (PI4P) (5). This study used TGN38/TGN46 as a marker for the TGN, yet TGN38/TGN46 cycles between the TGN and plasma membrane via endosomes (6). Thus, subsequent studies suggested that NLRP3 was recruited via PI4P to endosomal membranes in which TGN38/TGN46 became trapped (7, 8). Additional studies suggest NLRP3 may be recruited to the centrosome (9–11), and several other organellar membranes are also implicated in NLRP3 inflammasome activation (2). A critical limitation to-date, however, has been the dependence on ectopic overexpression in cell lines, and frequently, in cell lines that do not express inflammasome components, which has led to inconsistencies between studies of how and where NLRP3 inflammasomes form. Endogenous innate pathways are known to exhibit vastly different responses when compared to over-expression systems (12). Combined with the limited scope of existing reagents to visualise NLRP3, this has hampered the study of endogenous NLRP3 in primary cells, including understanding the nanoscale organisation of the endogenous NLRP3 inflammasome complex, and where and when NLRP3 contributes to inflammation *in vivo*.

Here, CRISPR/Cas9 was used to label endogenous NLRP3 in mice with the fluorescent reporter mScarlet-I. The data presented here in primary macrophages and in mice support a model where oligomeric NLRP3 associates with PI4P-positive membranes from multiple organelles, coordinating the oligomerisation of the ASC speck. Super resolution microscopy identified significant heterogeneity between the structures of NLRP3 and ASC within the speck, with nanoscopic differences in NLRP3 organisation observed in response to different stimuli, suggesting that the nanoscale structure of NLRP3 inflammasomes could be a critical regulator dictating inflammasome responses. *In vivo*, circulating monocytes and neutrophils are identified as the major cellular reservoirs of NLRP3 and are seen to migrate into tissues following a systemic inflammatory stimulus, highlighting monocytes and neutrophils as potential key cellular targets to limit NLRP3-driven diseases. Collectively, these paradigm-shifting insights highlight the NLRP3-mScarlet-I mouse as a powerful resource for the field.

## Results

### NLRP3-mScarlet-I is expressed and functional in BMDMs

CRISPR/Cas9 was used to edit the *Nlrp3* gene to generate an NLRP3 reporter mouse expressing an endogenous NLRP3 protein product with a C-terminal mScarlet-I tag, separated by a flexible linker to mitigate risk of perturbation of NLRP3 function (Fig 1a). Expression of wild type (WT) NLRP3 and NLRP3-mScarlet-I protein in bone marrow-derived macrophages (BMDMs) derived from WT mice and mice heterozygous and homozygous for the NLRP3-mScarlet-I gene was confirmed by western blot of cell lysates with anti-NLRP3 antibodies (Fig 1b,c). Priming with LPS increased NLRP3 protein levels as expected, and the levels of NLRP3 and NLRP3-mScarlet-I were comparable between genotypes. Although abundant bands were present at the expected molecular weights for tagged NLRP3 (∼145 kDa), a minor band (at ∼130 kDa) was also observed in NLRP3-mScarlet-I mice that may represent a truncated or cleaved form of the protein. Additionally, smaller bands at approximately 30 and 20 kDa were detected in the heterozygote and homozygote NLRP3-mScarlet-I cells with the anti-RFP/mScarlet-I antibody, consistent with minor mScarlet-I cleavage (Fig 1b,d,e). The temporal expression of NLRP3 and NLRP3-mScarlet-I were similar in response to different durations of LPS treatment (Fig 1f-h) and NLRP3 expression was also detected basally at comparable levels between WT and NLRP3-mScarlet-I BMDMs, suggesting the mScarlet-I tag does not impact normal protein turnover (Fig 1b-h). We confirmed NLRP3-mScarlet-I protein expression in peritoneal lavage cells directly isolated from WT, hetero- and homozygote NLRP3-mScarlet-I mice, which were cultured overnight and then primed with LPS (Supp Fig 1a-c). Endogenous NLRP3-mScarlet-I fluorescence in BMDMs was faint yet detectable by confocal microscopy (Fig 1i,j) but the signal could be amplified by immunofluorescence using anti-RFP/mScarlet-I labelling followed by Alexa Fluor 594-conjugated secondary antibodies (Fig 1k,l). Furthermore, LPS treatment increased the amount of NLRP3-mScarlet-I detected via the mScarlet-I tag (Fig 1i,j) and with signal amplification using anti-RFP labelling (Fig 1k,l), which corresponded to the increase in NLRP3-mScarlet-I protein detected via western blot (Fig 1b-h). In naïve and LPS-primed BMDMs, mScarlet-I signal was detected throughout the cell cytoplasm (Fig 1i,k). Higher confocal laser power was required to detect the non-amplified endogenous NLRP3-mScarlet-I signal compared with the amplified signal using the anti-RFP antibody, resulting in the detection of non-specific auto-fluorescent structures that were present in WT, heterozygote and homozygote NLRP3-mScarlet-I BMDMs (Fig 1i), likely reflecting auto-fluorescent lipofuscin within lysosomes, a trait associated with phagocytes (13).

**Figure 1.**
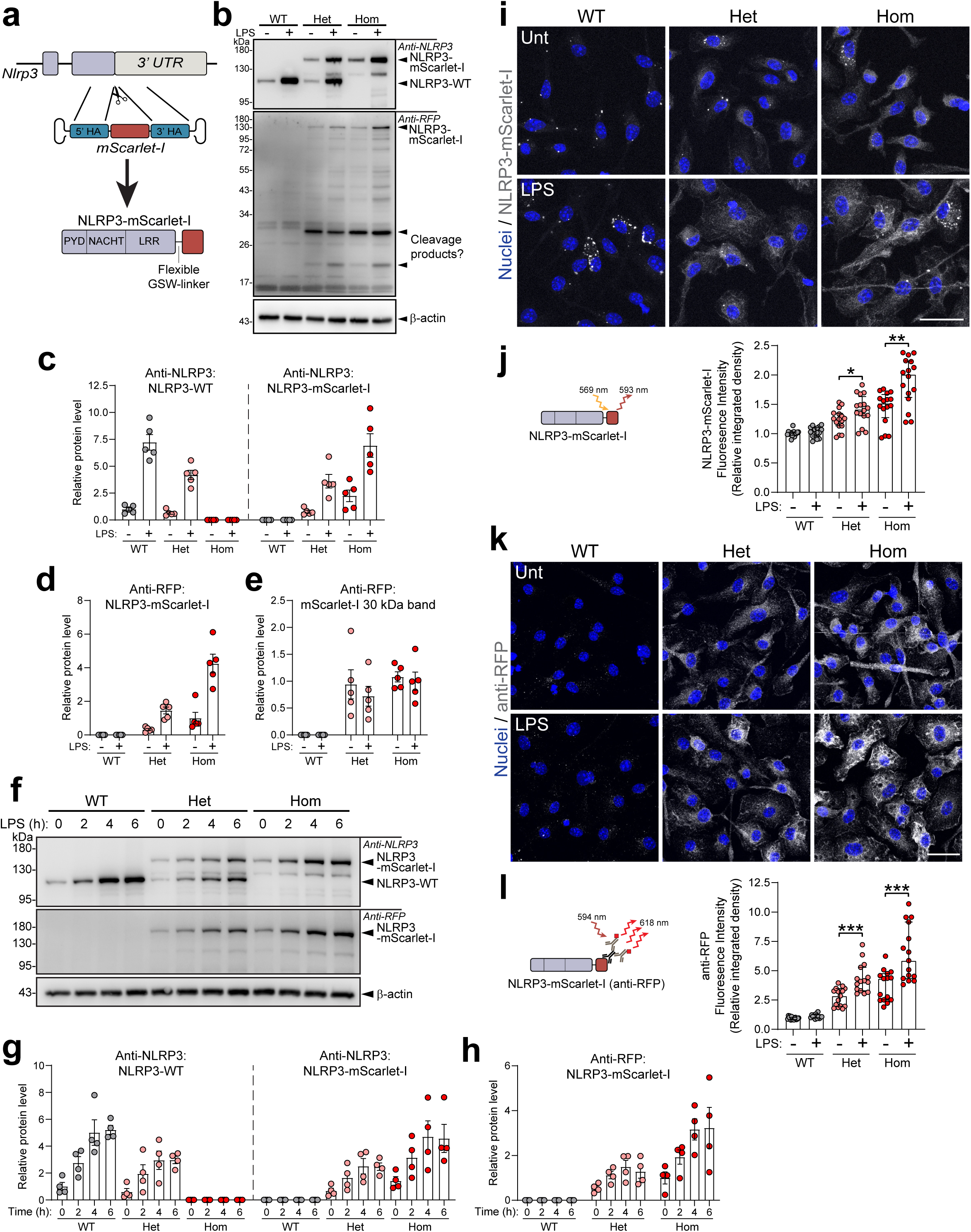
Endogenous NLRP3-mScarlet-I is expressed in primary mouse BMDMs. **(a)** Generation of the NLRP3-mScarlet-I reporter mouse. **(b-e)** BMDMs were untreated or primed with LPS (1 μg mL^-1^, 4 h) and (b) cell lysates were blotted for NLRP3 and mScarlet-I (representative from n=5). Densitometry of (c) anti-NLRP3 blots (NLRP3-WT (∼115 kDa) and NLRP3-mScarlet-I (∼145 kDa)) and (d,e) anti-RFP blots (NLRP3-mScarlet-I (∼145 kDa) and mScarlet-I 30 kDa band), normalised to β-actin (n=5). **(f-h)** BMDMs were primed with LPS (1 μg mL^-1^, 0-6 h) and (f) cell lysates were blotted for NLRP3 and mScarlet-I (representative from n=4). Densitometry of (g) anti-NLRP3 blots (NLRP3-WT and NLRP3-mScarlet-I) and (h) anti-RFP blots (NLRP3-mScarlet-I), normalised to β-actin (n=4). **(i-l)** BMDMs were untreated or primed with LPS (1 μg mL^-1^, 6 h), and NLRP3-mScarlet-I was detected by exciting the mScarlet-I tag (i, j) or with signal amplification by immunofluorescence using anti-RFP labelling (k, l; n=4; 3-5 FOV from 4 independent repeats). Maximum intensity projections are shown. Fluorescence intensity was measured on whole field of view. Scale bar is 25 μm. Data are mean ± SEM (c,d,e,g,h) or median ± IQR (j,l). Data were analysed using unpaired t-test (c, not significant, WT LPS (NLRP3-WT) vs Hom LPS (NLRP3-mScarlet-I); and l) or Mann-Whitney test (j). *p<0.05, **p<0.01, *** p<0.001.

To test whether NLRP3-mScarlet-I can form inflammasomes in response to activating stimuli, BMDMs isolated from WT, hetero-, and homozygous NLRP3-mScarlet-I mice were primed with LPS and stimulated with the NLRP3 inflammasome-activating stimuli nigericin, ATP, LLOME, imiquimod and silica, in the presence or absence of the NLRP3 specific inhibitor MCC950 (Fig 2a,b). These stimuli trigger NLRP3 activation through different mechanisms dependent upon ion flux (nigericin and ATP) (14), lysosome disruption (LLOME) (15), K^+^ efflux-independent routes (imiquimod) (16), and particulates causing frustrated phagocytosis (silica) (15). NLRP3 activation following exposure to intracellular LPS, which triggers activation of a non-canonical caspase-11 inflammasome pathway leading to NLRP3 inflammasome formation (17) was also assessed (Fig 2c,d). Activation of NLRP3 and NLRP3-mScarlet-I, as measured by release of IL-1β and by cell death, were comparable across stimuli except for imiquimod, where a genotype-dependent reduction in NLRP3-mScarlet-I activity was observed (Fig 2a,b). ASC oligomerisation blots confirmed that nigericin induced robust NLRP3 activation in both WT and NLRP3-mScarlet-I cells (Fig 2e), as well as cleavage and maturation of the downstream inflammasome proteins caspase-1, gasdermin D and IL-1β (Fig 2f). In response to imiquimod, ASC oligomerisation and maturation of caspase-1, gasdermin D and IL-1β were all reduced in NLRP3-mScarlet-I BMDMs, as compared to WT (Fig 2g,h). To further explore the apparent reduction in NLRP3 sensitivity to imiquimod in the NLRP3-mScarlet-I cells, two additional K^+^ efflux-independent activators, CL097 (16) and forchlorfenuron (18), were tested. Similarly to imiquimod, NLRP3-mScarlet-I activation in response to CL097 and forchlorfenuron was reduced compared to WT, as measured by IL-1β release and cell death (Supp Fig 2a,b). A comparison of dose responses of NLRP3 activation in WT and NLRP3-mScarlet-I BMDMs in response to imiquimod revealed a shift in the response curves, evidenced by a significant shift in EC_50_ values, although at higher doses of imiquimod the response of NLRP3-mScarlet-I was restored to a similar level to WT, as detected by IL-1β release and cell death (Fig 2i,j). A potential but non-significant decrease in LDH and IL-1β release was also observed at submaximal doses of nigericin in NLRP3-mScarlet-I cells (Fig 2k,l), but not with another K^+^ efflux-dependent stimulus silica (Fig 2m,n). Thus, these data suggest that NLRP3-mScarlet-I is functional with minor effects on sensitivity noted. To support these data, BMDMs were co-treated with imiquimod and monensin, previously demonstrated to potentiate the NLRP3 response to imiquimod (7). Co-treatment of imiquimod with monensin induced robust NLRP3-mScarlet-I activation at a dose of imiquimod which alone induced only a very weak response in NLRP3-mScarlet-I cells (Supp Fig 2c,d).

**Figure 2.**
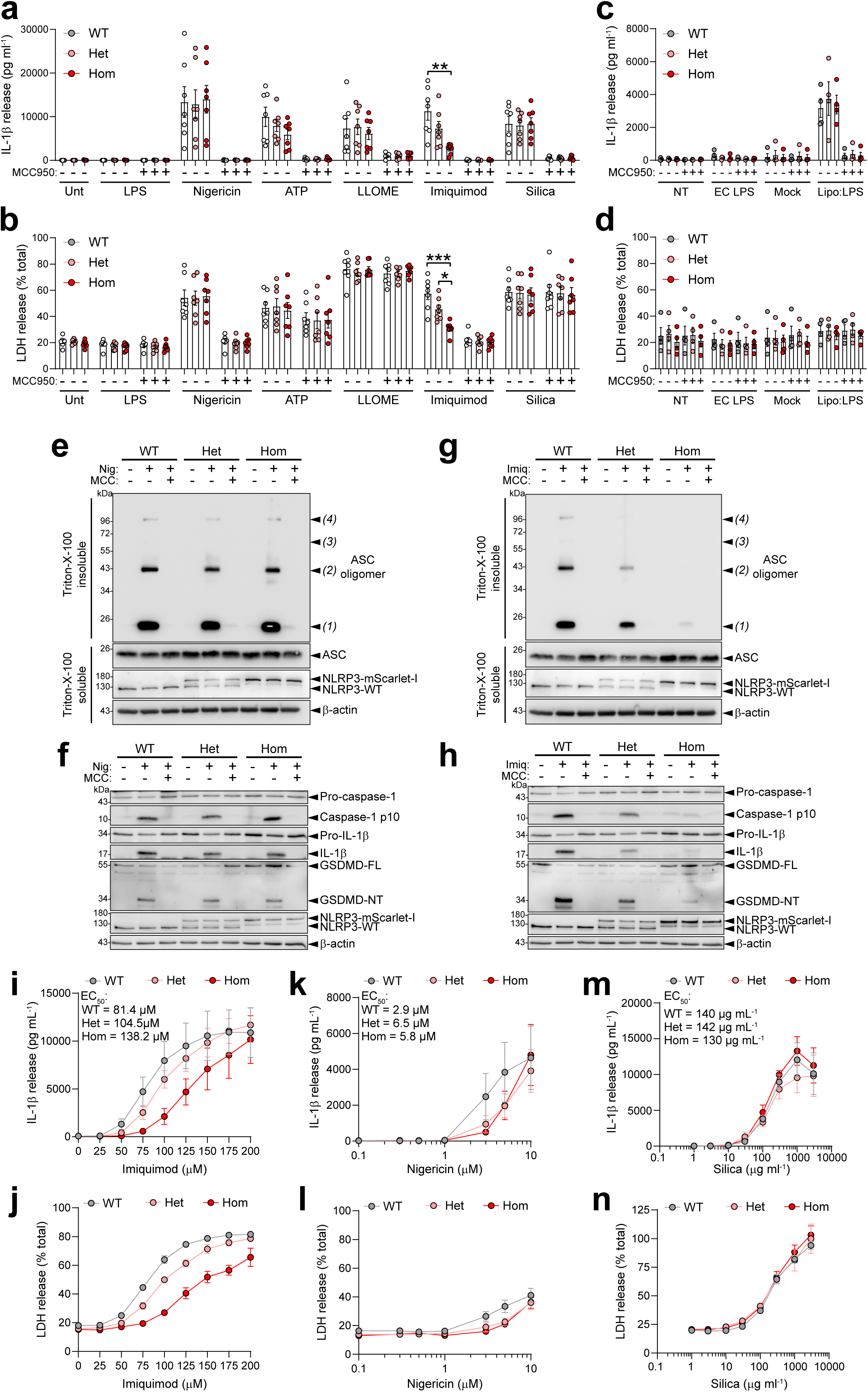
Endogenous NLRP3-mScarlet-I is active in primary BMDMs. **(a, b)** BMDMs were primed with LPS (1 μg mL^-1^, 4 h), and then treated ± MCC950 (10 μM, 15 min), before addition of nigericin (10 μM, 1 h), ATP (5 mM, 1 h), LLOME (1 mM, 1 h), imiquimod (75 μM, 2 h) or silica (300 μg mL^-1^, 2 h) (n=7). (a) IL-1β and (b) LDH release into the supernatant were measured. **(c-d)** BMDMs were primed with Pam3CSK4 (100 ng mL^-1^, 4 h) and then treated ± MCC950 (10 µM, 15 min), before addition of extracellular (EC) LPS (2 µg mL^-1^), lipofectamine 3000 reagent alone (mock) or LPS (2 µg mL^-1^, 20 h) complexed with lipofectamine 3000 (n=4). (c) IL-1β and (d) LDH release into the supernatant were measured. **(e-f)** BMDMs were primed with LPS (1 μg mL^-1^, 4 h) and then treated ± MCC950 (10 μM, 15 min), before addition of nigericin (10 μM, 1 h). Total cell lysates and supernatants were assessed for (e) DSS-crosslinked ASC oligomers (representative from n=4) and (f) cleavage of inflammasome proteins (representative from n=4). **(g-h)** BMDMs were primed with LPS (1 μg mL^-1^, 4 h) and then treated ± MCC950 (10 μM, 15 min), before addition of imiquimod (75 μM, 2 h). Total cell lysates and supernatants were assessed for (g) DSS-crosslinked ASC oligomers (representative from n=4) and (h) cleavage of inflammasome proteins (representative from n=4). **(i-n)** BMDMs were primed with LPS (1 μg mL^-1^, 4 h), and then treated with imiquimod (0-200 μM, 2 h), nigericin (0.1-10 μM, 1 h), or silica (1-3000 µg mL^-1^, 4 h). IL-1β release in response to (i) imiquimod, (k) nigericin and (m) silica. LDH release in response to (j) imiquimod, (l) nigericin and (n) silica. Data are mean ± SEM. Data were analysed using one-way ANOVA followed by Tukey’s post-hoc analysis (a-d). Concentration-response curves were fitted using a four-parameter logistical model to determine EC_50_ values (i, k, m). *p<0.05, **p<0.01, *** p<0.001.

### NLRP3-mScarlet-I is present within activated inflammasomes

To visualise NLRP3-mScarlet-I recruitment to the inflammasome complex during activation, WT and NLRP3-mScarlet-I BMDMs were treated with LPS and nigericin and then subsequently labelled with anti-RFP and anti-ASC antibodies. ASC speck formation was observed in response to nigericin in the WT, heterozygous and homozygous NLRP3-mScarlet-I BMDMs, and co-localised with an NLRP3-mScarlet-I speck in the heterozygous and homozygous cells (Fig 3a-c). MCC950 pre-treatment prevented the formation of both the NLRP3 and ASC specks, confirming that the inflammasome specks formed were NLRP3-dependent. A temporal analysis of NLRP3 inflammasome formation in response to nigericin revealed a time-dependent formation of NLRP3-mScarlet-I specks that colocalised with ASC specks (Supp Fig 3a-c). We could also observe endogenous NLRP3 specks in the absence of antibody amplification of the mScarlet-I immunofluorescence signal (Fig 3d-f). Consistent with earlier observations, auto-fluorescent puncta in the NLRP3-mScarlet-I channel in both WT and NLRP3-mScarlet-I BMDMs were occasionally detected, that did not colocalise with the ASC speck (Fig 3a,d). Thus, both endogenous NLRP3-mScarlet-I and ASC were visualised within the same NLRP3 inflammasome complex. In BMDMs heterozygous for NLRP3-mScarlet-I, the proportion of NLRP3-mScarlet-I-positive ASC specks was the same as cells homozygous for NLRP3-mScarlet-I, suggesting that in the heterozygous cells, NLRP3-mScarlet-I molecules are incorporated into the inflammasome complex alongside WT NLRP3 molecules. The presence of NLRP3 molecules in the Triton-soluble and -insoluble fractions as a proxy for NLRP3 oligomerisation was also assessed, and following nigericin stimulation, enriched WT NLRP3 and NLRP3-mScarlet-I molecules were detected in the Triton-insoluble fraction, alongside ASC (Fig 3g). Thus, these data suggest that NLRP3-mScarlet-I molecules are in the active inflammasome complex, meaning NLRP3-mScarlet-I is an ideal tool to investigate formation and structure of NLRP3 inflammasome complexes.

**Figure 3.**
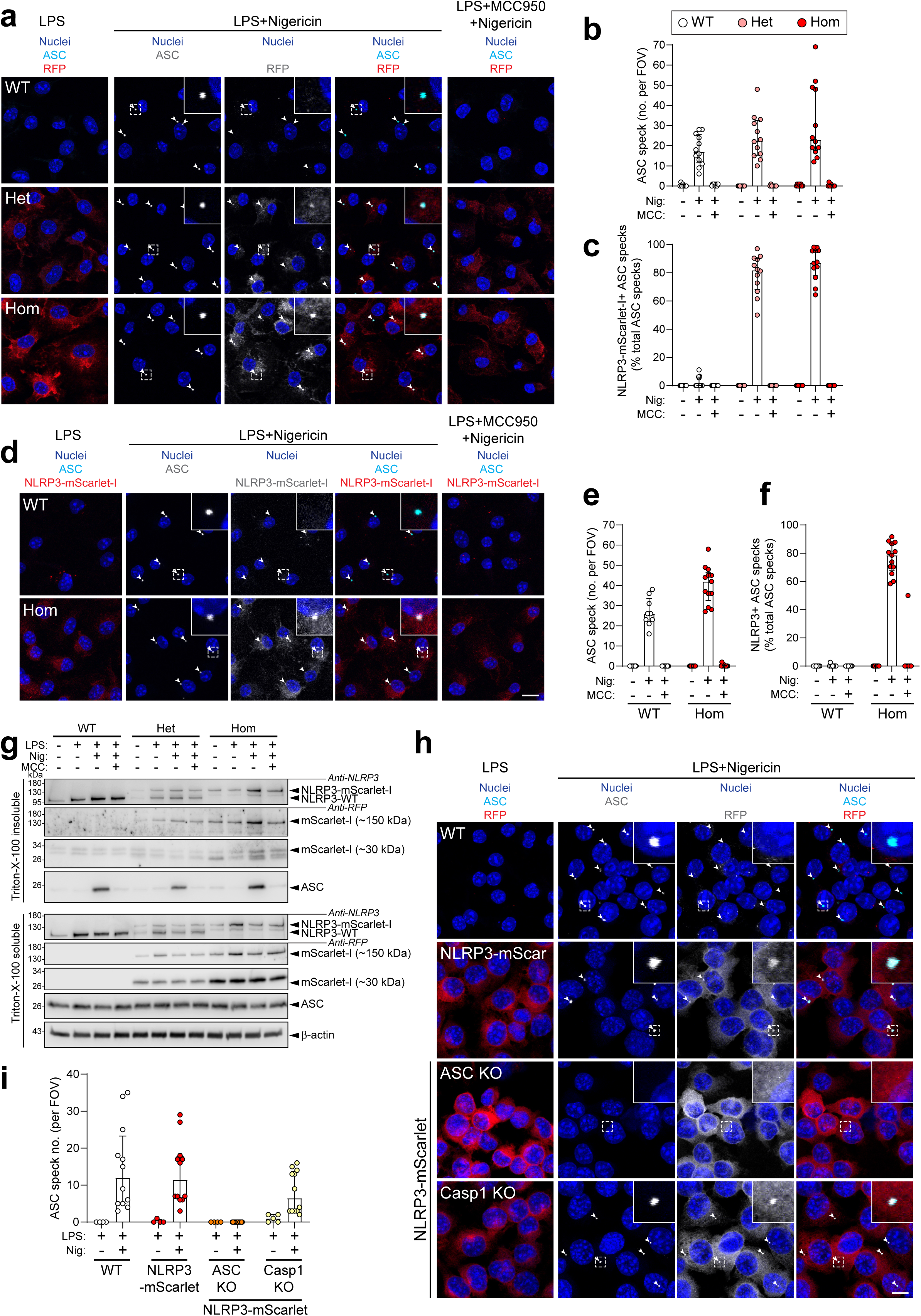
NLRP3-mScarlet-I is present within activated inflammasomes. **(a-c)** BMDMs were primed with LPS (1 μg mL^-1^, 6 h) and then treated with VX-765 (10 μM, 15 min, to prevent pyroptosis) ± MCC950 (10 μM, 15 min), before addition of nigericin (10 μM, 1 h) (n=3 – 2-4 FOV from 3 biological repeats). (a) Cells were assessed by immunofluorescence for ASC (green) and NLRP3-mScarlet-I using anti-RFP labelling of mScarlet-I (red). A single z-plane is shown. Scale bar = 20 μm. (b) ASC speck number per FOV and (c) % of NLRP3-mScarlet-I+ ASC specks were quantified. **(d-f)** BMDMs were primed with LPS (1 μg mL^-1^, 6 h) and then treated with VX-765 (10 μM, 15 min, to prevent pyroptosis) ± MCC950 (10 μM, 15 min), before addition of nigericin (10 μM, 1 h) (n=3 – 2-4 FOV from 3 biological repeats). (d) Cells were assessed by immunofluorescence for ASC (green) and endogenous NLRP3-mScarlet-I signal (red). Maximum intensity projections are shown. Scale bar = 10 μm. (e) ASC speck number per FOV and (f) % of NLRP3-mScarlet-I+ ASC specks. **(g)** Triton X-100-soluble and -insoluble fractions were generated from WT, NLRP3-mScarlet-I heterozygote or NLRP3-mScarlet-I homozygote BMDMs, treated ± LPS (1 µg mL^-1^, 4 h) and then treated ± MCC950 (10 μM, 15 min), before addition of nigericin (10 μM). Fractions were blotted for NLRP3, ASC and RFP (mScarlet-I) (representative from n=4). **(h,i)** WT, NLRP3-mScarlet-I, ASC KO NLRP3-mScarlet-I, or Casp1 KO NLRP3-mScarlet-I iBMDMs were treated with LPS (1 µg mL^-1^, 4 h) and then treated with VX-765 (10 µM, 15 min) before the addition of nigericin (10 µM, 1 h). (h) Cells were assessed by immunofluorescence for ASC (green) and NLRP3-mScarlet-I using anti-RFP labelling of mScarlet-I (red). Maximum intensity projections are shown. Scale bar = 10 μm (n=4, 3-5 FOV from 4 passages). (i) ASC speck number per FOV was quantified. Data are median ± IQR.

Previous reports have demonstrated that in the absence of the adaptor protein ASC, NLRP3 forms multiple puncta spread throughout the cell, rather than a single large speck (5, 7, 19). To assess this with NLRP3-mScarlet-I, immortalised BMDMs (iBMDMs) from homozygote NLRP3-mScarlet-I BMDMs were generated and CRISPR-Cas9 used to knockout ASC or caspase-1. ASC and caspase-1 KO NLRP3-mScarlet-I iBMDMs were confirmed to be selectively deficient in ASC and caspase-1 respectively (Supp Fig 4a) and lacked NLRP3 inflammasome responses, demonstrated by an absence of IL-1β and LDH release to nigericin treatment (Supp Figure 4b,c). Following nigericin treatment, both ASC and NLRP3 specks were detected in NLRP3-mScarlet-I and caspase-1 KO NLRP3-mScarlet-I iBMDMs, as in the primary cells, but in the ASC KO NLRP3-mScarlet-I iBMDMs, the loss of ASC completely prevented formation of the NLRP3 speck (Fig 3h,i), suggesting that the oligomerisation and localisation of NLRP3 and ASC are inter-dependent. Notably, multiple NLRP3 puncta were not observed in the absence of ASC. Whilst unable to form an inflammasome, ASC KO NLRP3-mScarlet-I iBMDM cells retained nigericin-induced signalling upstream of the inflammasome, exhibited by endolysosomal trafficking disruption in response to nigericin, detected through disruption of perinuclear TGN38 signal (Supp Fig 4d).

### Sub-cellular localisation of endogenous NLRP3 inflammasomes in primary macrophages

The sub-cellular localisation of NLRP3 at endogenous levels in primary macrophages remains unclear. To resolve the sub-cellular location of endogenous inflammasome complexes, we activated WT and NLRP3-mScarlet-I primary BMDMs, labelled them for NLRP3-mScarlet-I, ASC, and different organelles or the lipid PI4P, and analysed using confocal microscopy. Markers used were for PI4P (Fig 4a, Supp Fig 5a), the early endosome marker EEA1 (Fig 4b, Supp Fig 5b), the lysosome marker LAMP1 (Fig 4c, Supp Fig 5c), the TGN marker TGN38 (Fig 4d, Supp Fig 5d) and the centrosome marker γ-tubulin (Fig 4e, Supp Fig 5e). Caspase-1-dependent cleavage of markers, and pyroptosis, were prevented by the inclusion of the caspase-1/11 inhibitor VX-765 in all experiments. In LPS-treated BMDMs, PI4P was detected in the perinuclear region, likely reflecting its presence at the TGN. Nigericin increased PI4P signal intensity on punctate structures throughout the cell (Fig 4a, Supp Fig 5a), suggesting an increase in PI4P levels at other cellular compartments, consistent with previous reports (8). Analysis of confocal z-stacks was conducted to observe whether any direct associations between PI4P and the NLRP3/ASC specks were present. Direct associations were determined by either overlapping signal or immediate contact within the XY- or Z-planes, within the resolution permitted by confocal microscopy. Based on this analysis, 76±5% of ASC and 74±5% of NLRP3 specks showed association with PI4P following nigericin treatment (Fig 4f,g, Supp Fig 5f). This analysis was applied to the other organelle markers used. Significant associations were observed between the inflammasome and the early endosome marker EEA1, the lysosome marker LAMP1, and the TGN marker TGN38 (Fig 4b-d, Supp Fig 5b-d). There was marked variability in association with markers, with 79±5% of NLRP3 specks overlapping or in immediate contact with LAMP1, 30±2% with TGN38, and 37±4% with EEA1 vesicles (Fig 4f,g, Supp Fig 5f). Similar levels of association were observed with the ASC speck, with 77±4% of ASC specks associated with LAMP1, 34±4% with TGN38, and 38±6% with EEA1 vesicles (Fig 4f,g, Supp Fig 5f). These associations either reflect the ability of PI4P-containing membranes from multiple organelles to interact with the NLRP3/ASC speck, or that confocal microscopy is not sufficient to resolve true associations over very close proximities caused by crowding in the peri-nuclear space. There were limited interactions of NLRP3/ASC specks observed with γ-tubulin, a centrosome marker, in response to nigericin, with only 8±3% of NLRP3 specks associated with γ-tubulin puncta (Fig 4e-g, Supp Fig 5e,f). The association of NLRP3/ASC specks with γ-tubulin increased to 33±4% in response to imiquimod, consistent with a recent study (20), although the majority of NLRP3/ASC specks did not colocalize with the centrosome, regardless of stimulus (Fig 4e-g). Given that the centrosome consists of 1-2 clear puncta, it was feasible to measure the shortest distance between the ASC speck and the centrosome in the XYZ plane. ASC specks formed after imiquimod stimulation were on average in closer proximity to the centrosome compared to those formed following nigericin stimulation (Fig 4h). These data support a model in which endogenous NLRP3 interacts with PI4P-positive membrane across multiple organelles, and to a lesser extent the centrosome, to co-ordinate inflammasome assembly, and suggest that the subcellular site of inflammasome assembly varies between NLRP3 stimuli.

**Figure 4.**
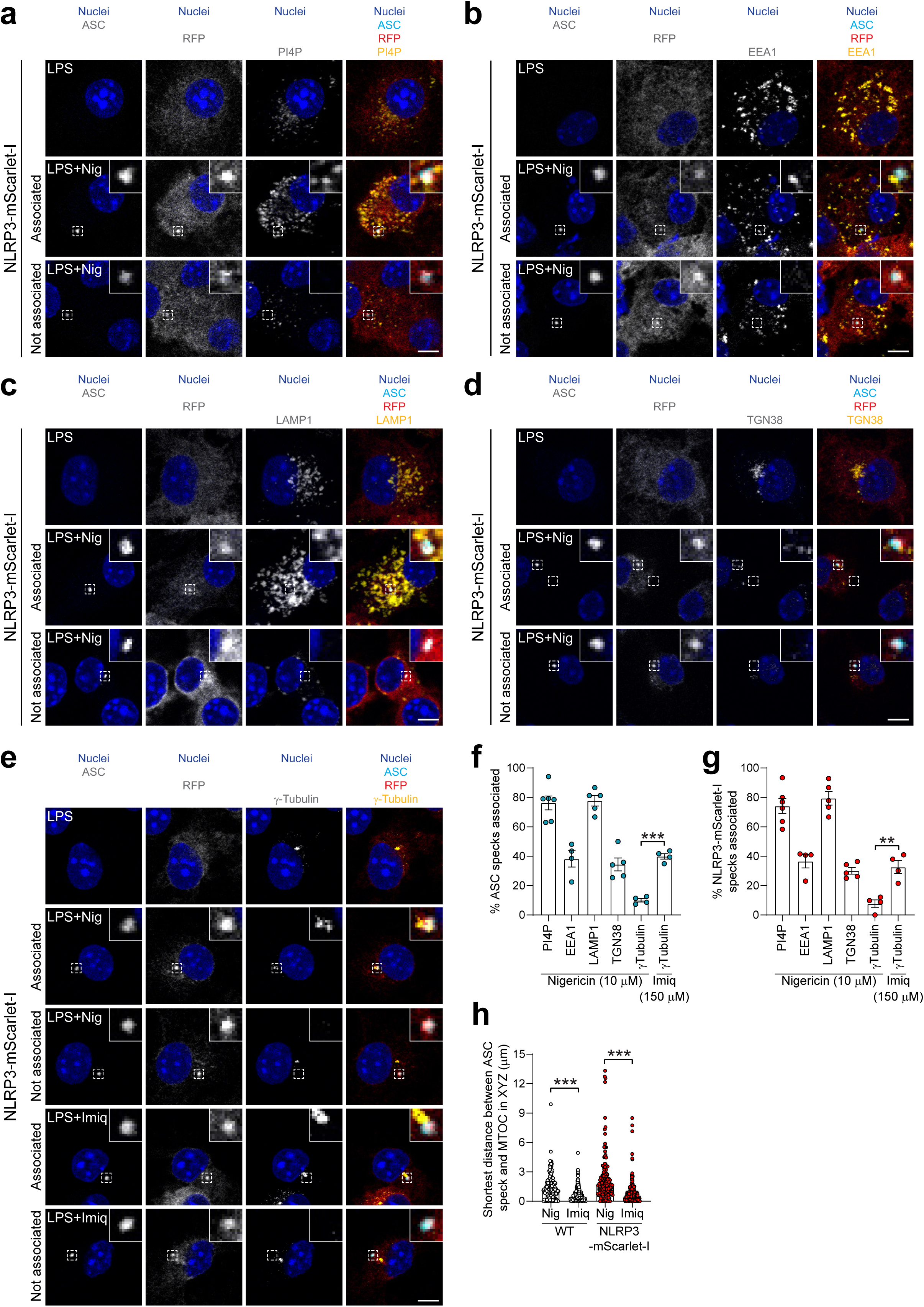
Sub-cellular localisation of endogenous NLRP3 inflammasomes in primary macrophages. Primary NLRP3-mScarlet-I BMDMs were primed with LPS (1 μg mL^-1^, 6 h) and then treated with VX-765 (10 μM, 15 min, to prevent pyroptosis), before addition of nigericin (10 μM, 30 min) or imiquimod (150 μM, 30 min). **(a-e)** Cells were assessed by immunofluorescence for ASC (green) and NLRP3-mScarlet-I using anti-RFP labelling of mScarlet-I (red), as well as (a) PI4P, (b) EEA1, (c) LAMP1, (d) TGN38, or (e) γ-tubulin (yellow) (n=4-6 – 3-4 FOV averaged from 4-6 biological repeats). Example NLRP3/ASC specks that were classified as ‘associated’ or ‘not associated’ with the respective organelle marker are shown. Single z-planes are shown. Scale bar = 5 μm. **(f-g)** Association of (f) ASC specks and (g) NLRP3-mScarlet-I specks with each organelle marker, based on all specks from that biological repeat. **(h)** Shortest distance of every ASC speck from the MTOC (closest γ-tubulin punctum) in the XYZ plane in WT and NLRP3-mScarlet BMDMs was manually determined. Data are mean ± SEM (f,g) or median ± IQR (h). Data were analysed using unpaired t-test (f,g) or Kruskal-Wallis test with Dunn’s multiple comparisons post-hoc analysis (h). *p<0.05, **p<0.01, *** p<0.001.

### Nanoscale NLRP3 inflammasome speck structure is heterogeneous and stimulus-dependent

The nanostructure of ASC within the speck was recently reported to consist of a dense core surrounded by a less dense periphery containing filament-like structures using direct stochastic optical reconstruction microscopy (dSTORM) (21). Super-resolution microscopy has been performed for endogenous NLRP3 using primary and secondary antibodies (22), although antibody amplification and accessibility to the dense speck complex may have limited nanoscale resolution (21). The NLRP3-mScarlet-I mice allowed us to use an mScarlet-I nanobody to directly label NLRP3-mScarlet-I. Thus, we could visualise tagged endogenous NLRP3 within inflammasome complexes in primary cells. In combination with a previously described nanobody to ASC (23) and by using dSTORM, we were able to compare the nanoscopic architecture of the ASC and NLRP3 specks. Consistent with previously published data (21), dSTORM images showed that ASC specks within nigericin- and imiquimod-treated LPS-primed WT and NLRP3-mScarlet-I primary BMDMs have a dense core surrounded by a less dense and filamentous periphery (Fig 5a, Supp Fig 6-8). NLRP3-mScarlet-I-positive signal co-localised with, and was distributed across, the ASC speck (Fig 5a). Thresholding the signal of NLRP3-mScarlet-I and ASC permitted quantification of NLRP3 and ASC area within the inflammasome complex. There were no significant differences in ASC speck size between WT and NLRP3-mScarlet-I BMDMs, suggesting addition of the mScarlet-I tag to endogenous NLRP3 did not impact the ASC speck nanostructure (Fig 5b). Interestingly, stimulus-specific differences were observed in the structure of the inflammasome complexes. Imiquimod-induced ASC and NLRP3-mScarlet-I specks were greater in area compared to those formed by nigericin treatment, suggesting that differences in signalling pathways caused by the different stimuli influence inflammasome structure and composition (Fig 5b,c). A weak, but significant, correlation between NLRP3-mScarlet-I and ASC speck area in both nigericin- and imiquimod-induced specks was observed, suggesting that the number of ASC molecules within specks is partially dependent on the number of NLRP3 molecules (Fig 5d,e). Comparing the ratio of NLRP3-mScarlet-I/ASC area within the inflammasome speck also revealed large variability in the composition of ASC and NLRP3 in specks between cells, with some specks containing up to twice as much staining area for ASC versus NLRP3 and vice versa (Fig 5f). This variability may partially reflect the stochastic nature of dSTORM imaging, rather than the true speck structure. We also examined roundness of NLRP3-mScarlet-I and ASC in the inflammasome speck, where a value of 1 represents a perfect circle which decreases with increasing angularity. Both NLRP3-mScarlet-I and ASC specks had similar roundness values of ∼0.75 and this wasn’t affected by genotype or stimulus (Supp Fig 9a,b). A weak, but significant, correlation comparing NLRP3-mScarlet-I and ASC roundness suggested a relationship exists between the two (Supp Fig 9c,d).

**Figure 5.**
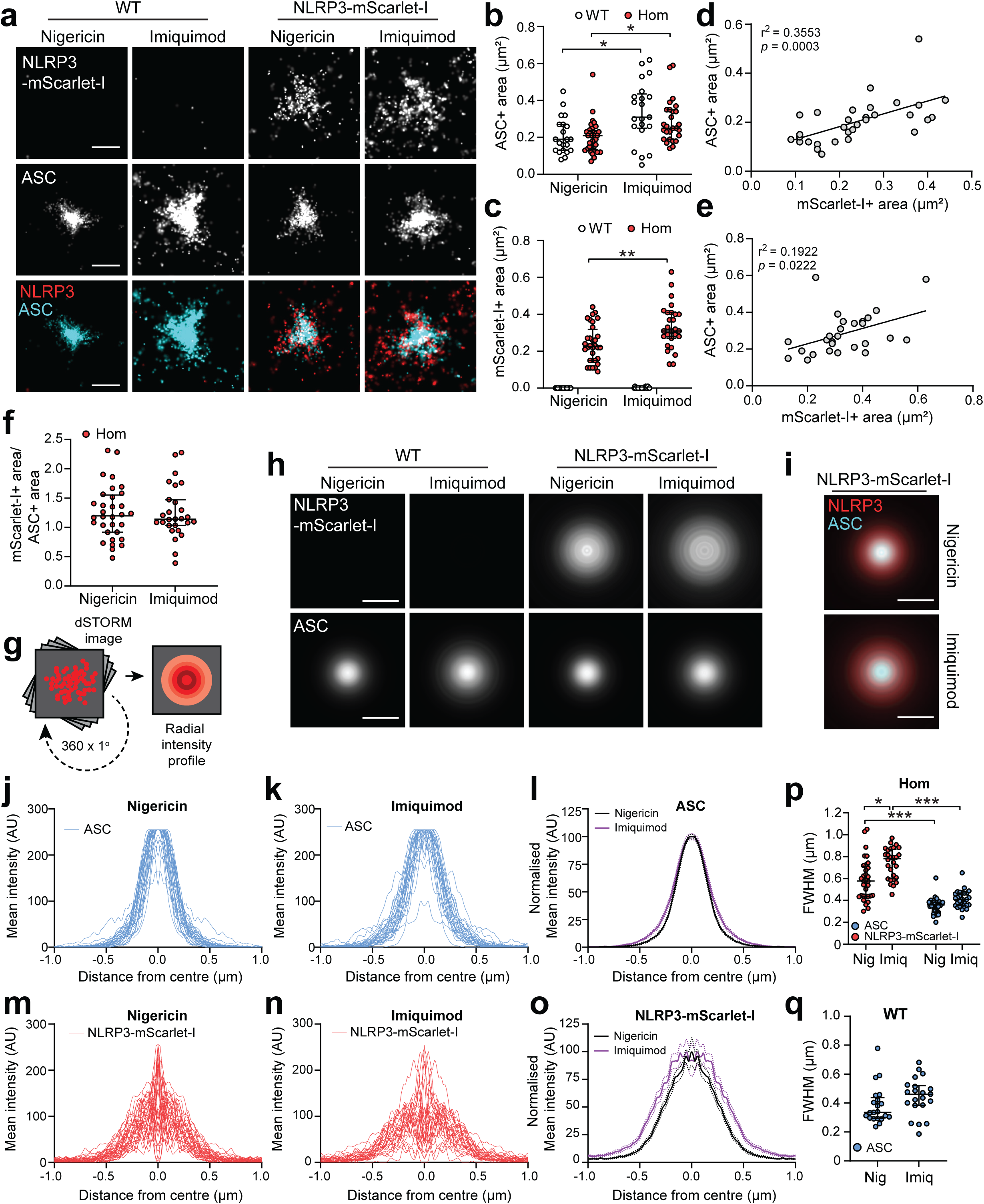
Nanoscale NLRP3 inflammasome speck structure is heterogeneous and stimulus-dependent. BMDMs were primed with LPS (1 μg mL^-1^, 6 h) and then treated with VX-765 (10 μM, 15 min), before addition of nigericin (10 μM, 1 h) or imiquimod (150 μM, 1 h) (n=5 biological repeats, 4-11 specks imaged per biological repeat). **(a)** Representative two-colour dSTORM images of NLRP3-mScarlet-I and ASC within nanobody-labelled inflammasome complexes (scale bar = 0.5 µm). **(b)** Area of ASC-positive and **(c)** NLRP3-mScarlet-I-positive staining in inflammasome specks. **(d,e)** Correlations of ASC and NLRP3-mScarlet-I-positive area of staining in (d) nigericin- and (e) imiquimod-induced specks. **(f)** Quantification of the ratio of NLRP3-mScarlet-I:ASC-positive area within inflammasome specks. **(g)** Schematic of generating radial intensity profiles from dSTORM images. **(h)** Radial averages generated from NLRP3-mScarlet-I and ASC specks dSTORM images. Averages generated from following number of specks: WT, nigericin 21, imiquimod 21; NLRP3-mScarlet-I Homozygous, nigericin 32, Imiquimod 28 (scale bar = 0.5 µm). **(i)** Merged NLRP3-mScarlet-I and ASC radial averages from (g) (scale bar = 0.5 µm). **(j-l)** Individual (j) nigericin, (k) imiquimod and (l) mode-normalised average radial intensity profiles of ASC within NLRP3-mScarlet-I BMDMs. **(m-o)** Individual (m) nigericin, (n) imiquimod and (o) mode-normalised average radial intensity profiles of NLRP3-mScarlet-I within specks of NLRP3-mScarlet-I BMDMs. **(p-q)** Quantification of full width at half maximum (FWHM) of Gaussian fits in homozygous NLRP3-mScarlet-I BMDMs (p) and WT BMDMs (q) of NLRP3-mScarlet-I and ASC speck radial intensity profiles. Data are median ± IQR. Data were analysed using Kruskal-Wallis test with Dunn’s multiple comparisons post-hoc analysis (b,p) or unpaired t-test (c,f,q). Correlations were determined using a simple linear regression. *p<0.05, **p<0.01, *** p<0.001.

Radial averaging analysis (24) on dSTORM images of inflammasome complexes was used to assess the generalised distribution of NLRP3-mScarlet-I and ASC molecules in relation to the central point of the inflammasome complex (Fig 5g). By collating radial intensity plots from every NLRP3 and ASC speck captured in primary NLRP3-mScarlet-I BMDMs in response to nigericin (Supp Fig 6) and imiquimod (Supp Fig 7) it was possible to generate an averaged radial intensity profile plot for NLRP3 and ASC within the inflammasome complex (Fig 5h-o; Supp Fig 6-7). The same analysis was conducted for ASC in primary WT BMDMs (Supp Fig 8). This further demonstrated the differences between the ASC and NLRP3 distribution, with ASC forming a dense core whilst NLRP3 was distributed more diffusely over a larger area with higher variability. To quantify the distribution of NLRP3 and ASC signal in relation to the central point of the inflammasome complex, the full width at half maximum (FWHM) of the Gaussian fits of individual NLRP3-mScarlet-I and ASC radial intensity profiles in WT and NLRP3-mScarlet-I cells were calculated (Fig 5p,q, Supp Fig 9e-g). Consistent with our visual observations, NLRP3 exhibited a significantly broader distribution than ASC in both nigericin- and imiquimod-induced inflammasomes. Furthermore, NLRP3-mScarlet-I was distributed over a significantly larger area following imiquimod treatment compared with nigericin (Fig 5p). Importantly, the distribution of ASC was identical between NLRP3-mScarlet-I and WT BMDMs, showing the ASC speck formed normally (Supp Fig 9h-i). Together, these data reveal a previously unappreciated marked heterogeneity in inflammasome structure and highlight stimulus-dependent differences in the organisation of NLRP3 in the speck, but that the overall organisation of ASC was not significantly different when studied at this resolution.

### Circulating and recruited myeloid cells express NLRP3 in vivo

*In vivo* levels of endogenous NLRP3 protein, at the single cell level, at homeostasis and during inflammation has yet to be demonstrated. We used spectral flow cytometry to measure NLRP3 protein abundance across immune cells in the blood, bone marrow and spleen, three major immune-relevant tissues, from NLRP3-mScarlet-I mice. We also assessed NLRP3 protein expression in the brain and dura, given the importance of NLRP3 in neuroinflammation and brain disease (25) (Fig 6a-e). By thresholding against the signal detected in cells from WT littermate mice, we determined both the percentage of NLRP3-mScarlet-I-expressing cells (Fig 6f-j) and, by examining the geometric mean fluorescence intensity (gMFI) of NLRP3-mScarlet-I per cell population, their relative levels of NLRP3 protein (Fig 6k-o). NLRP3-mScarlet-I protein was detected in the innate immune cell compartment, predominantly in monocytes, neutrophils and conventional dendritic cells (cDC)2, and at lower levels in resident macrophage, microglia and cDC1 populations (Fig 6). The relatively high abundance of NLRP3-mScarlet-I in monocytes, neutrophils and cDC2 in naïve mice was surprising, given that induction of NLRP3 by proinflammatory NF-κB signalling is typically considered a prerequisite for NLRP3 inflammasome activation (1). NLRP3-mScarlet-I signal was not detected, or detected at very low levels, in NK cells, B-, or T-cells across all tissues tested (Fig 6). These data show that the NLRP3 protein is widely expressed in homeostatic conditions *in vivo*, particularly in circulating monocytes and neutrophils.

**Figure 6.**
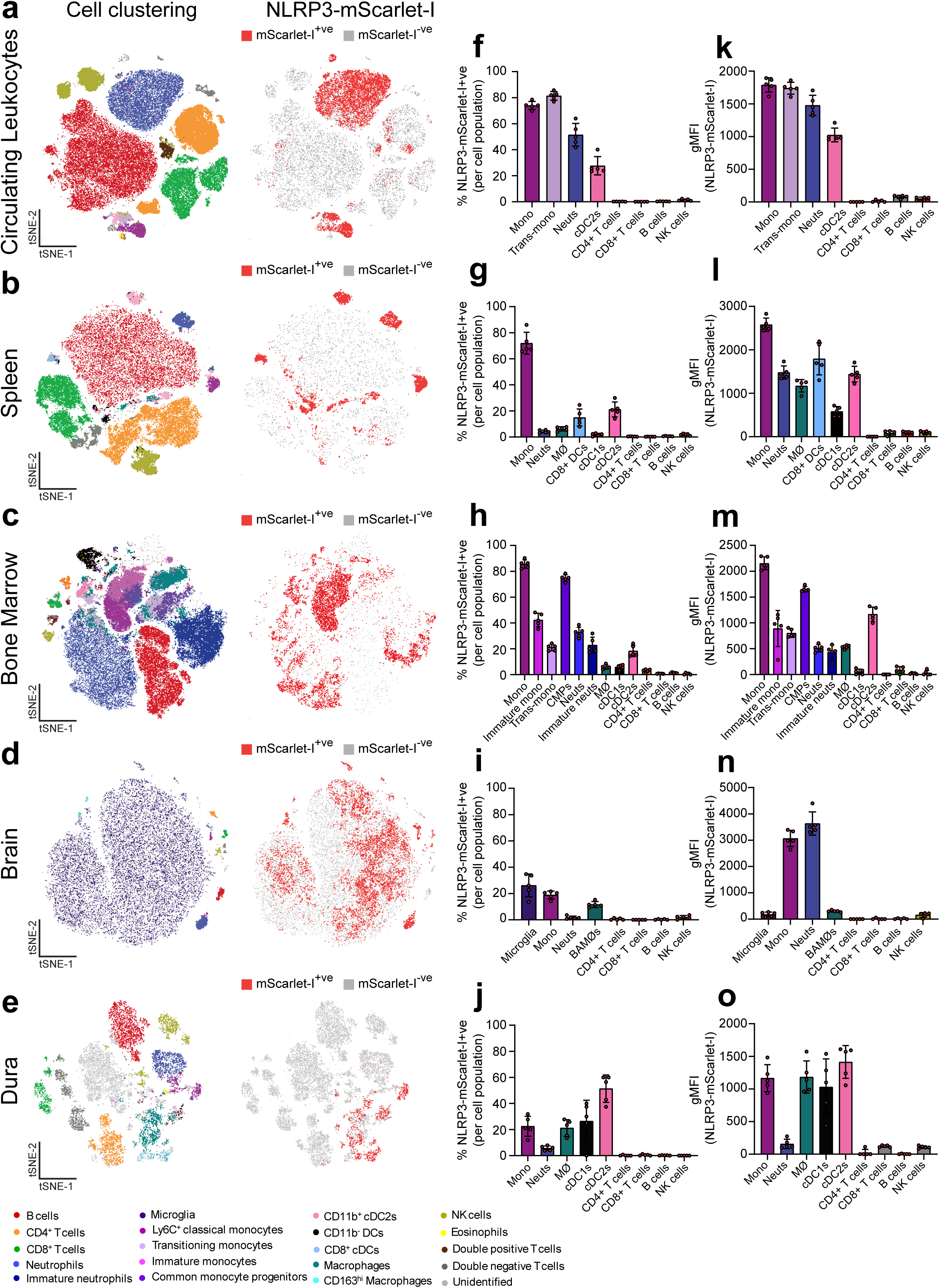
Circulating myeloid cells are the predominant source of NLRP3 at homeostasis *in vivo*. **(a-e)** tSNE plots from spectral flow cytometry analysis of NLRP3-mScarlet-I expression in CD45-positive immune cells isolated from circulating leukocytes (a), spleen (b), bone marrow (c), brain (d) and dura (e) from naïve homozygous NLRP3-mScarlet-I mice (generated from single cells from n=5). **(f-j)** Percentage of NLRP3-mScarlet-I positive cells in each indicated cell population in circulating leukocytes (f), spleen (g), bone marrow (h), brain (i) and dura (j) (n=5). **(k-o)** Geometric mean fluorescence (gMFI) of NLRP3-mScarlet-I in circulating leukocytes (f), spleen (g), bone marrow (h), brain (i) and dura (j) (n=5). gMFI was corrected to fluorescence measured in WT mice to control for autofluorescence. Abbreviations: Mono, *monocytes*; Trans-mono, *transitioning monocytes*; Neuts, *neutrophils*; cDC, *conventional dendritic cell*; MØ, *macrophage*; CMP, *common myeloid progenitor*; BAMØ, *border-associated macrophage (CD163^hi^)*.

Next, to examine the expression profile of NLRP3 in inflammation and to determine whether NLRP3-mScarlet-I is activated normally *in vivo*, an LPS-induced peritonitis model was used where IL-1β processing is known to be NLRP3-dependent (26, 27). WT, hetero- and homozygote mice for NLRP3-mScarlet-I were injected i.p. with LPS (10 mg kg^-1^) and 4 h later the peritoneal lavage, blood, brain and dura were collected. In all mice, LPS induced an increase in IL-1β, IL-18, and IL-6 levels in plasma and peritoneal lavage, and this was comparable between genotypes (Fig 7a-e), showing that the mScarlet-I tag does not interfere with NLRP3 inflammasome function *in vivo*. At the immune cell level, cells in the peritoneum, blood, brain and dura were assessed for NLRP3 protein abundance following LPS (Fig 7f-i, Supp Fig 10) and the levels were quantified (Fig 7j-m). The proportion of neutrophils and resident macrophages expressing NLRP3-mScarlet-I protein, and its abundance in these cells, increased in response to LPS in the peritoneum, blood, brain and dura (Fig 7f-m). Monocytes, on the other hand, exhibited a varied response, with a decrease in the gMFI and the percentage of monocytes positive for NLRP3-mScarlet-I in the peritoneum (Fig 7f,j, Supp Fig 10a,e), the site of LPS injection. Since this model induces NLRP3 inflammasome activation in the peritoneum (Fig 7a-e) (27), the reduction in monocytic NLRP3-mScarlet-I could be due to NLRP3-expressing cells undergoing pyroptosis. Similarly, NLRP3-mScarlet-I-positive monocytes, peritoneal macrophages, and neutrophils, were all reduced as a proportion of the total number of CD45-positive cells in the peritoneum (Fig 7f, Supp Fig 10a,e). However, in response to LPS, we observed an increase in the percentage of monocytes in the brain and dura positive for NLRP3-mScarlet-I (Fig 7h,i, Supp Fig 10c,d) and a corresponding increase in gMFI (Fig 7l,m). In the brain and surrounding dura, NLRP3-mScarlet-I-positive monocytes significantly increased as a percentage of the total CD45-positive cells following LPS treatment, as well as NLRP3-mScarlet-I-positive neutrophils in the dura, suggesting that circulating leukocytes with high NLRP3 levels were starting to infiltrate into tissues distal to the site of LPS injection. Of note, there was no significant increase in NLRP3-mScarlet-I-positive microglia detected (Supp Fig 10c,g), with only a slight increase in NLRP3-mScarlet-I gMFI relative to other cell populations following LPS treatment (Fig 7l). Whilst there was a slight decrease in the gMFI and percentage of NLRP3-mScarlet-I-positive cDC1s in the peritoneum (Fig 7j, Supp Fig 10a,e), NLRP3-mScarlet-I abundance in cDC1 and cDC2 cells was unchanged in response to LPS across all tissues tested (Fig 7j-m, Supp Fig 10), despite already expressing relatively high levels of NLRP3 in homeostatic conditions (Fig 6). In the predominantly NLRP3-mScarlet-I-negative populations (T cells, B cells, NK cells), there was no induction of NLRP3 following LPS (Fig 7j-m, Supp Fig 10). Together, these studies reveal striking cell- and tissue-specific differences in NLRP3 expression *in vivo* at homeostasis and during acute inflammation. Specifically, these data reveal that alongside resident macrophages, monocytes and neutrophils are the predominant NLRP3-expressing cell types that are rapidly recruited into tissues during acute systemic inflammation. Therefore, the extent and location of NLRP3-driven inflammation may be determined as much by the recruitment of NLRP3-high leukocytes as by local tissue responses.

**Figure 7.**
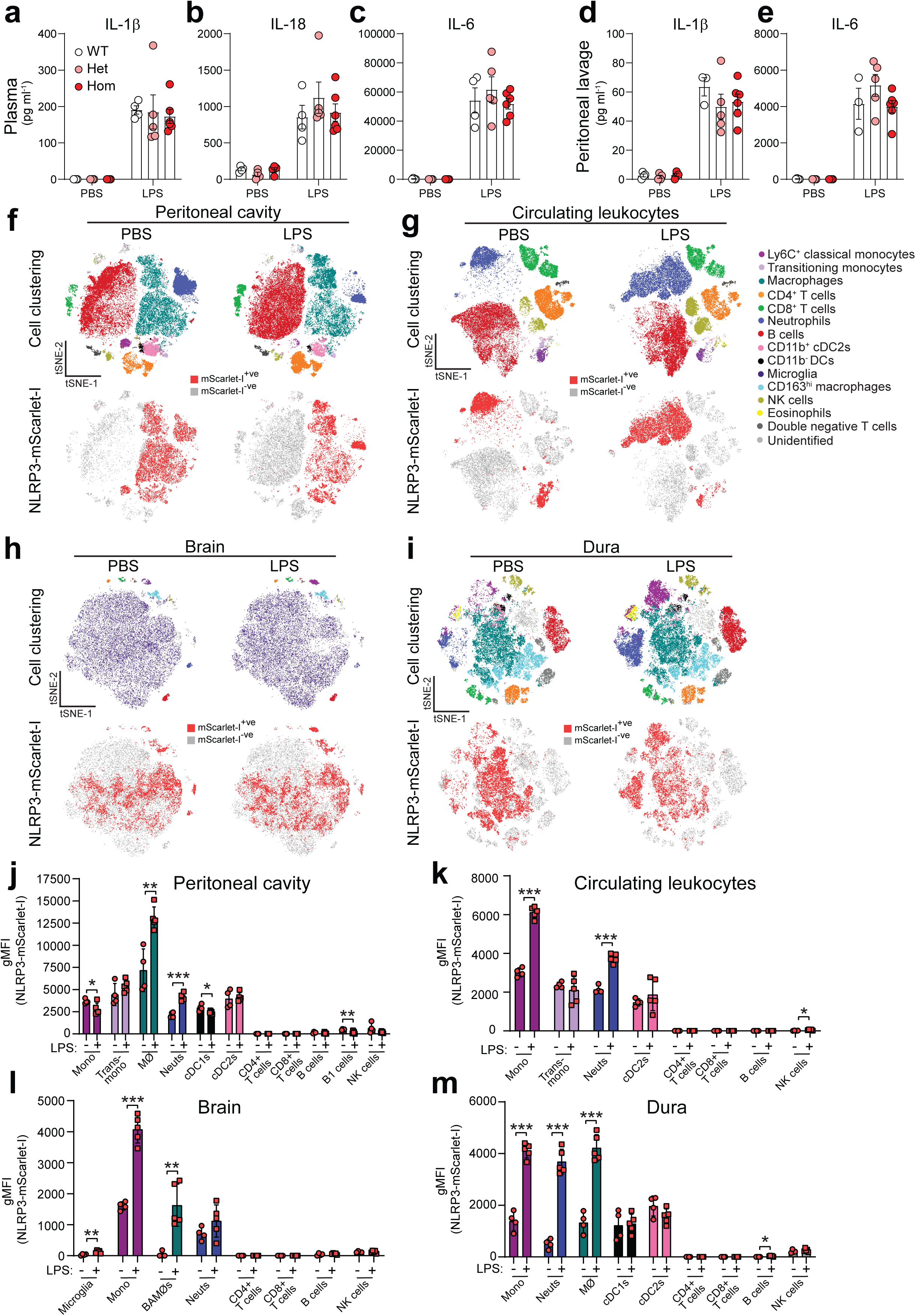
NLRP3 expression and activation in response to LPS-induced peritonitis. WT, NLRP3-mScarlet-I heterozygotes or homozygotes were treated with intraperitoneal LPS (10 mg kg^-1^, 4 h) or PBS control. **(a-c)** ELISA on plasma for IL-1β (a), IL-18 (b), and IL-6 (c) (n=4-6). **(d-e)** ELISA on peritoneal lavage for IL-1β (d) and IL-6 (e) (n=3-6). **(f-i)** tSNE plots from spectral flow cytometry analysis of NLRP3-mScarlet-I expression in immune cells isolated from the peritoneum (f), circulating leukocytes (g), brain (h) and dura (i) from homozygous NLRP3-mScarlet-I mice treated with intraperitoneal PBS (n=4) or LPS (10 mg kg^-1^, 4 h) (n=5). **(j-m)** gMFI of NLRP3-mScarlet-I expression in indicated cell populations in the peritoneum (j), circulating leukocytes (k), brain (l) and dura (m) (n=4-5). gMFI was corrected to fluorescence measured in WT mice to control for autofluorescence. Data are mean ± SEM. Data were analysed using Kruskal-Wallis test with Dunn’s multiple comparisons (a,b) or one-way ANOVA followed by Tukey’s post-hoc analysis (c-e), or an unpaired t-test (j-m, PBS vs LPS per cell type). *p<0.05, **p<0.01, *** p<0.001. Abbreviations: Mono, *monocytes*; Trans-mono, *transitioning monocytes*; Neuts, *neutrophils*; cDC, *conventional dendritic cell*; MØ, *macrophage*; BAMØ, *border-associated macrophage (CD163^hi^)*.

Our data suggest that circulating NLRP3-high monocytes and neutrophils may be recruited to tissues during inflammation to co-ordinate rapid inflammasome responses. This may be particularly relevant in the brain, with relatively low abundance of NLRP3 within resident microglia, but high levels in perivascular macrophages and recruited monocytes and neutrophils following LPS administration (Fig 7l,m). Therefore, to further understand the dynamics and cellular localisation of NLRP3-mScarlet-I within the brain during acute inflammation, cranial windows were inserted and real-time intravital imaging of NLRP3-mScarlet-I in the brain was performed during systemic inflammation induced by LPS-peritonitis. WT or NLRP3-mScarlet-I homozygous mice were treated with LPS (i.p. 10 mg kg^-1^, 6 hours), and NLRP3-mScarlet-I fluorescence was examined using confocal microscopy (Fig 8a). We imaged at 561 nm to visualise mScarlet-I and at 488 nm to identify non-specific autofluorescence obtained due to the higher laser powers required for visualising endogenous NLRP3-mScarlet-I. NLRP3-mScarlet-I-positive cells were localised within the vasculature in NLRP3-mScarlet-I mice, which was absent in WT mice (Fig 8b, Supp Video 1). Our flow cytometry analysis (Fig 7g,k Supp Fig 10b) suggested that these cells were predominantly monocytes and neutrophils. Supporting this, NLRP3-mScarlet-I-positive cells adhered to, and rolled along, the endothelium in NLRP3-mScarlet-I mice (Fig 8c, Supp Video 1), features characteristic of leukocytes in inflamed conditions (28). Administration of anti-CD45-FITC, a pan-immune cell marker, revealed leukocytes adhering and rolling to the endothelium in WT and NLRP3-mScarlet-I mice, which co-localised with NLRP3-mScarlet-I (Supp Fig 11, Supp Video 1). Interestingly, NLRP3-mScarlet-I signal above background fluorescence was not detectable outside the blood vessels in the brain parenchyma (Fig 8b,c), matching our flow cytometry data (Fig 7l). Therefore, to explore if the NLRP3-mScarlet-I cells that adhered and rolled on the vasculature were ultimately recruited into the brain, we performed an extended LPS-peritonitis model (two doses of 1 mg kg^-1^ LPS i.p., 12 h apart), as was previously shown to cause significant leukocyte recruitment into the brain (29, 30), and visualised NLRP3-mScarlet-I signal in the brain at 12 and 24 hours after the first LPS injection (Fig 8d). At 12 hours post LPS, we observed NLRP3-mScarlet-I-positive cells arrested to the side of vessels, a key step preceding diapedesis, but the NLRP3-mScarlet-I signal was still predominantly restricted to cells within the vasculature (Fig 8e, Supp Video 2). However, 24 hours post LPS, there was a reduction in NLRP3-mScarlet-I-positive cells adhered to vessels, accompanied by the appearance of multiple extravascular NLRP3-mScarlet-I positive cells, predominantly located in the perivascular space (Fig 8f, Supp Video 2). Extravascular NLRP3-mScarlet-I-positive cells were highly motile and could be observed migrating along vessels and through the parenchyma (Fig 8g, Supp Video 2). Some of these extravascular NLRP3-mScarlet-I-positive cells exhibited prominent lipofuscin/lysosomal autofluorescence moving within the cell, suggesting that these are phagocytes, such as monocyte-derived macrophages (Fig 8f, Supp Video 2). Thus, we demonstrate for the first time real-time intravital visualisation of NLRP3 expressed at endogenous levels *in vivo,* capturing the recruitment and infiltration of NLRP3-high leukocytes into the brain during acute systemic inflammation. Together, our findings reveal infiltrating monocytes and neutrophils as a major source of NLRP3 during acute neuroinflammation, revealing an underappreciated contribution of peripheral leukocytes in driving inflammasome responses in the brain.

**Figure 8.**
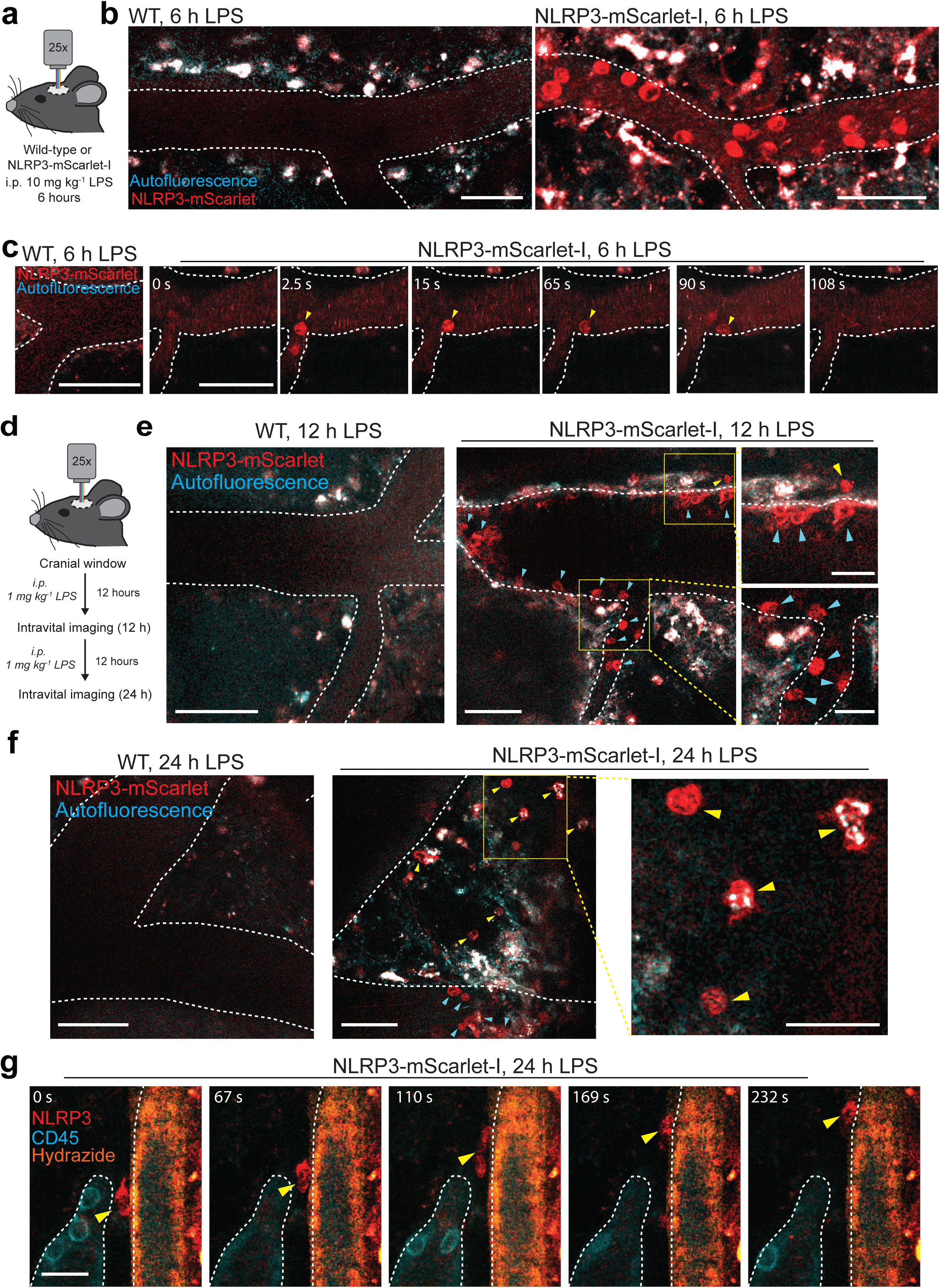
Infiltration of circulating NLRP3-mScarlet-I-positive cells into the brain during systemic inflammation. **(a)** Intravital imaging of the brain was performed in wild-type (WT) and NLRP3-mScarlet-I littermates following insertion of cranial windows and intraperitoneal (i.p.) injection of LPS (10 mg kg^-1^, 6 h). **(b)** Representative confocal image of NLRP3-mScarlet-I (561 nm, red) and auto-fluorescent background (488 nm, cyan), with resulting non-specific autofluorescence labelled as white. The vasculature is shown by the dotted white line. Scale bar = 50 µm (n=4). **(c)** Timelapse from experiments in (a) of NLRP3-mScarlet-I-positive cells rolling along the vasculature. A representative rolling cell is tracked by a yellow arrow. Time shown in seconds (s), Scale bar = 50 µm (n=4). **(d)** Intravital imaging of the brain in WT and NLRP3-mScarlet-I littermates following insertion of cranial windows and i.p. injection of LPS (1 mg kg^-1^, administered twice, 12 h apart), imaged at 12 h and 24 h. **(e-f)** Representative images at 12 h (e) and 24 h (f) post first LPS injection. The vasculature is shown by the dotted white line. Intravascular NLRP3-mScarlet-I-positive cells are indicated by blue arrows and extravascular NLRP3-mScarlet-I-positive cells are indicated by yellow arrows. Scale bar = 50 µm, scale bar in inset = 25 µm. (n=3 NLRP3-mScarlet-I, n=2 wild-type). **(g)** Timelapse from experiments in (f) of a representative extravascular NLRP3-mScarlet-I-positive cell migrating along the perivascular space. NLRP3-mScarlet-I (red), intravenously administered anti-CD45-FITC (cyan), and hydrazide (yellow, labels elastin in artery walls). Scale bar = 20 µm. (n=3).

## Discussion

This study describes the generation and characterisation of a mouse where the endogenous *Nlrp3* gene was tagged with the fluorescent protein mScarlet-I, facilitating study of NLRP3 at endogenous levels in primary cells and *in vivo* leading to leaps in our understanding of NLRP3 biology. A question that has emerged within the literature is the nature of the organelle upon which NLRP3 associates for its activation. Previous research from this group (7) and others (8) proposed endosomes and lysosomes as sites for NLRP3 nucleation, while vesicles of TGN are also proposed (5) amongst other organelles (2), in addition to the non-membranous centrosome (9, 10). Supporting previous observations, data presented here suggest the lipid PI4P is strongly associated with the majority of endogenous NLRP3 inflammasome complexes. PI4P, which is produced by phosphatidylinositol-4-kinases (PI4K), is a major regulator of Golgi function (31) and is also present on endosomal, lysosomal (32) and plasma membranes (33). PI4P accumulates on endosomal and lysosomal membranes following disruption of ER-endosomal contact sites and lysosomal damage, respectively (8, 34, 35), both features that commonly occur following treatment with NLRP3 activators (8, 34–36). Inhibition of PI4Ks that regulate endolysosomal PI4P pools, but not PI4Ks that maintain the Golgi PI4P pool, prevents NLRP3 recruitment to endosomes (8). In addition to disrupting endosomal cargo trafficking (7, 8), NLRP3-activating stimuli also cause widespread proteomic changes within organelles (37), highlighting the general stress exerted by NLRP3-activating stimuli, and that PI4P on the membranes of stressed organelles is likely key to NLRP3 sensing. Recent work shows that ectopic expression of NLRP3 specifically targeted to different organelles results in inflammasome formation (38). It is thus possible that any PI4P-containing membrane could provide a scaffold for NLRP3 inflammasome activation. Given the size and structure of the NLRP3 speck observed, it is also conceivable that even within the same cell, the NLRP3 speck could be coordinated by multiple nucleation points from different organelle membranes. Limited colocalisation of NLRP3 specks with the centrosome were observed in the present study in response to nigericin, although specks formed by imiquimod stimulation showed a greater association and were closer to the centrosome. These differences may be due to a milder effect of imiquimod on trafficking and subcellular reorganisation compared to other NLRP3-activating stimuli (7, 37), or potentially due to a difference in the mechanism of activation (20). Correlative cryo-light and cryo-electron microscopy (CLEM) have also been used to visualise the NLRP3 inflammasome complex and surrounding cellular environment (39, 40). Supporting observations presented here, Liu *et al* found nigericin-induced inflammasome complexes form a tubular core with branched filaments in the proximity of Golgi- and endosome-like vesicles that were often near, but physically separated from, the MTOC (within 2-3 µm) in mouse iBMDMs (39). Conversely, Wang *et al* found using over-expressed NLRP3-mScarlet in human NLRP3-KO THP-1, that NLRP3 puncta localise with the MTOC (40). The discrepancy between these studies, in addition to species differences, could potentially be due to protein expression levels. Supporting this point, our previous data shows that over-expressed NLRP3 in THP-1 forms two puncta, one localised at the ASC speck and one at the MTOC (11).

Using nanobodies to mScarlet-I and ASC, this study has revealed the nanoscale structure of NLRP3 and ASC within inflammasome complexes in primary macrophages expressed at endogenous levels. This confirmed previously published data showing the ASC speck consists of a dense core and a less dense, filamentous periphery (21), and provides new data on the organisation of the NLRP3 speck. While these data show colocalisation of NLRP3-mScarlet-I and ASC in the same speck, they reveal different patterns of organisation. These data also reveal that the NLRP3-mScarlet-I speck formed in response to imiquimod is larger than the NLRP3-mScarlet-I speck formed in response to nigericin – again highlighting differences in the mechanisms driving inflammasome formation, or in the sub-cellular organisation of the cell in response to different stimuli. The formation of the NLRP3 speck was also found to be entirely dependent upon ASC, as ASC KO NLRP3-mScarlet-I iBMDMs did not form NLRP3 specks. These data suggest the NLRP3 and ASC specks are completely co-dependent upon each other for their formation. Thus, it may be suggested that as oligomerising NLRP3 on PI4P membranes is pulled together by the oligomerising ASC, NLRP3 is detached from some of the nucleating membranes into the speck-like structure. In this scenario, the variable organisation of the NLRP3 speck may reflect the variable distribution of organellar membranes providing a scaffold.

The NLRP3-mScarlet-I mouse also provides new insights into the abundance of NLRP3 protein in different cell types *in vivo*. Single cell protein abundance of NLRP3 across tissues has not yet been reported. We reveal that in naïve mice, in blood, the main cellular reservoirs of NLRP3 are monocytes and neutrophils, suggesting that *in vivo* these are the rapid responders that can drive NLRP3 inflammasome responses to threats. Conversely, in brain tissue, NLRP3 protein is expressed in resident microglia, but in fewer cells and at much lower levels than circulating monocytes and neutrophils. Previous research from this group has reported recruitment of IL-1-expressing monocytes and neutrophils into the brain parenchyma in models of both ischaemic stroke (41) and intracerebral haemorrhage (ICH) (42). A recent study reports that increased leukocyte NLRP3 in blood correlates with worse stroke outcome (43). Sustained exposure to systemic inflammatory stimuli can also promote monocyte and neutrophil recruitment to the brain (29, 30). Our real-time intravital imaging revealed that circulating NLRP3-expressing leukocytes infiltrate the brain in acute inflammatory conditions, uncovering a previously underappreciated mechanism by which recruitment of peripheral immune cells may shape the magnitude of NLRP3-driven inflammation within the CNS. As such, targeting NLRP3 activity in the brain could be possible by targeting peripheral myeloid cells before they are recruited, potentially bypassing the need for a drug to directly cross the blood-brain barrier, thus paving the way for new therapeutic targeting strategies for brain diseases. Beyond the brain, recruitment of NLRP3-expressing leukocytes may also be a critical step underpinning aberrant inflammasome responses in tissues across the body, redefining our approaches to target NLRP3 across a range of inflammasome-driven pathologies.

In conclusion, generation of the NLRP3-mScarlet-I mouse has revealed new insight into the regulation and organisation of the NLRP3 inflammasome expressed at endogenous levels in primary cells, and has provided further insights into NLRP3 responses *in vivo*. This study highlights the importance of endogenous gene tagging and provides a valuable resource to study NLRP3 without the artefacts caused by overexpression. Together, these findings establish the NLRP3-mScarlet-I mouse as a powerful platform for studying inflammasome biology across health and disease. By enabling visualisation of NLRP3 from the molecular scale to whole tissues *in vivo*, this model will facilitate mechanistic studies into endogenous inflammasome regulation, NLRP3-driven pathologies, and the cellular sources of inflammasome activity within disease, to further evaluate the potential of NLRP3 as a therapeutic target across a broad range of inflammatory disorders.

## Methods

### Generation of Nlrp3–mScarlet-I knock-in mice

NLRP3-mScarlet-I mice were generated and maintained at the University of Manchester. A C-terminally tagged *Nlrp3*–mScarlet-I knock-in mouse line was generated using CRISPR/Cas9–mediated HDR. To mitigate for risk of perturbation on NLRP3 function, a flexible Gly and Ser rich linker, GGSGGSGSAGSAAGSGEF (termed GSWaldo or GSW, modified from Waldo *et al*, (1999) (44) was used. Two sgRNAs targeting sequences proximal to the *Nlrp3* stop codon were designed (sg588: 5′-TCGAGATTTTCCTGGTAGGCG-3′; sg590: 5′- GTCCTGCTTCCACGCCTACC-3′) and synthesised as full length sgRNA (Integrated DNA Technologies, Coralville, USA). The donor construct consisted of ∼400bp homology arms to the targeted region flanking the GSW linker followed by the mScarlet-I fluorescent protein, to be fused in-frame at the C terminus of *Nlrp3*. The final allele sequence is provided (Supp Fig 12).

HDR donor DNA was delivered using an adeno-associated virus (AAV)–based system (serotype 6: Vectorbuilder) following a modified CRISPR + AAV donor approach (45). One-cell C57BL/6J embryos were thawed and incubated in 25µL EmbryoMax^®^ KSOM Mouse Embryo Media, (Merck Sigma-Aldrich, product: MR-121) with 25µL AAV donor (≥2 × 10^11^ genome copies mL⁻¹) in an atmosphere of mixed gas (5% CO_2_, 5% O_2_) at 37°C for 4 h before electroporation. sgRNAs were complexed with recombinant Cas9 protein (NEB) to form ribonucleoproteins (RNPs). Transduced zygotes were electroporated with RNPs using a Nepagene system (CUY501P1-1.5 electrode; 4 poring pulses at 40 V, 3.5 ms; 5 transfer pulses at 5 V, 50 ms (46) and transferred at the one-cell stage into pseudopregnant CD-1(ICR) females.

Following birth of pups, genomic DNA harvested from ear biopsies (Sigma REDExtract-N-Amp Tissue PCR kit) was screened by PCR using primer pairs spanning the integration site. A locus-specific primer pair, which lie outside the homology arms (DB35_gF: 5′-CACTGCCCAGCCCAATCTG-3′; DB35_gR3: 5′-GGCTGCCACAAACCTTCCAT-3′) was used to assess on-target editing, and knock-in–specific amplification was performed using a combination of genomic and insert-specific primers (mScarlet-I_F: 5′-CGCGTGATGAACTTCGAGGA-3′; mScarlet-I_R: 5′-CTTGTACAGCTCGTCCATGCC-3′). Candidate knock-in alleles were identified by amplification of correctly sized HDR products and validated by Sanger sequencing of PCR amplicons generated with high-fidelity polymerase. Sequencing confirmed precise in-frame integration of the GSW–mScarlet-I tag at the *Nlrp3* C terminus. Truncated alleles arising from incomplete HDR were excluded from further analysis. A correctly targeted founder was crossed to wild-type C57BL/6J mice. Germline transmission was confirmed in F1 offspring by PCR and sequencing across the integration site. Multiple independent F1 animals carrying the correctly targeted allele were obtained and used to establish the colony on a C57BL/6J background (*C57BL/6J.Nlrp3*^em1Jagre^).

Animals were housed in ventilated cages with temperature and humidity maintained between 20–24°C and 45-65%, respectively, with a 12 h light-dark cycle. All procedures were performed with appropriate personal and project licenses in place, in accordance with the Home Office (Animals) Scientific Procedures Act (1986), approved by the Home Office and the local Animal Ethical Review Group, University of Manchester, and reported according to the ARRIVE guidelines.

### Cell culture

Primary bone marrow-derived macrophages (BMDMs) were generated using bone marrow isolated from femurs and tibias from wild-type, NLRP3-mScarlet-I hetero-and homozygote littermates. Isolated bone marrow was subjected to red blood cell lysis, passed through a 70 µm cell strainer, and cultured in DMEM (10% v/v FBS, 100 U mL^-1^ penicillin, 100 µg mL^-1^ streptomycin) supplemented with L929-conditioned media (30 % v/v) for 6-7 d. BMDM cultures were fed on day 3-4 with extra media containing L929-conditioned media. Prior to experiments, BMDMs were scraped and seeded overnight at a density of 1 x 10^6^ mL^-1^ (unless stated otherwise) in DMEM (10% v/v FBS, 100 U mL^-1^ penicillin, 100 µg mL^-1^ streptomycin).

Peritoneal macrophages were isolated via peritoneal lavage. Following cervical dislocation, mice were injected intraperitoneally with 6 mL RPMI (3% v/v FBS, 1 mM EDTA) and the lavage was collected. Peritoneal cells were centrifuged at 500 *x g* for 5 min before resuspension in DMEM (10% v/v FBS, 100 U mL^-1^ penicillin, 100 µg mL^-1^ streptomycin) and plated overnight at a density of 1 x 10^6^ mL^-1^.

Immortalised BMDMs (iBMDMs) were cultured in DMEM (10% v/v FBS, 100 U mL^-1^ penicillin, 100 µg mL^-1^ streptomycin). Prior to experiments, iBMDMs were scraped and seeded overnight at a density of 0.75 x 10^6^ mL^-1^.

### Generation, immortalisation, and clonal selection of NLRP3-mScarlet-I iBMDMs

Immortalisation of NLRP3-mScarlet-I BMDMs was performed using a previously published protocol (47). Bone marrow was isolated from the femurs and tibias of NLRP3-mScarlet-I mice. Macrophage differentiation was induced by culturing the cells in complete DMEM supplemented with L929-conditioned medium (20% v/v) at 37°C in a humidified 5% CO_2_ atmosphere. To generate the retrovirus, Cre-J2 producer cells were grown to 90% confluency, at which point the media was replaced. After 24 h, the retrovirus-containing supernatant was harvested, passed through a 0.45 μm filter to remove cellular debris, and stored at -80°C. Three days post-isolation, primary macrophage progenitors were transduced by replacing the culture media with a 1:1 mixture of fresh L929-conditioned medium and the Cre-J2 retroviral supernatant. A second identical round of viral infection was performed 48 h later. Seven days following the second infection, the selection of iBMDMs was initiated by reducing the L929-conditioned media supplementation to 10%. Over a subsequent period of 2 to 6 months, the concentration of L929-conditioned media was progressively halved every 1 to 2 weeks. This gradual withdrawal of exogenous growth factors (M-CSF) led to the death of non-infected primary macrophages, allowing only the successfully immortalised cells to survive and eventually proliferate in the complete absence of L929 supplementation. To validate the macrophage phenotype and establish monoclonal cell lines, the fully immortalised bulk cells were harvested, washed, and prepared for fluorescence-activated cell sorting (FACS). Cells were stained with a panel of fluorophore-conjugated antibodies targeting murine macrophage surface markers: CD45-BV510, CD11b-PeCy7, and F4/80-BV785. DAPI (UV excitation) was included to assess cell viability. Using FACS, single viable (DAPI-negative) cells that exhibited a robust CD45^+^ CD11b^+^ F4/80^+^ signature, while retaining the endogenous NLRP3-mScarlet-I reporter expression, were sorted directly into 96-well plates at a density of 1 cell per well. The resulting monoclonal colonies were subsequently expanded and validated for downstream functional assays.

### Pycard and Casp1 gene knockout in iBMDMs from Nlrp3-mScarlet-I mouse

Single-guide RNAs (sgRNAs) were designed to target the first asymmetric exon shared by all annotated isoforms of *Pycard* (ASC) or *Casp1*. Candidate sgRNAs were identified using the CRISPR Finder tool on the Wellcome Sanger Institute Genome Editing (WGE) platform. Two sgRNAs per gene were selected based on predicted on-target efficiency and minimal off-target potential. The following sgRNA sequences (5′–3′) were used: *Pycard* (ASC) sgRNA945: CTATCTGGAGTCGTATGGCT and sgRNA947: CAAACTTGTCAGCTACTATC, *Casp1* sgRNA346: CGAGTGGTTGTATTCATTAT, sgRNA347:GAGGGCAAGACGTGTACGAG. For all experiments, paired sgRNAs targeting the same gene were delivered simultaneously as ribonucleoprotein (RNP) complexes. Recombinant *Streptococcus pyogenes* Cas9 nuclease (Alt-R S.p. Cas9 Nuclease V3; Integrated DNA Technologies) was combined with synthetic sgRNAs at a molar ratio of 2:1 (sgRNA:Cas9).

For cell transfection, we developed an optimised CRISPR KO protocol in iBMDMs as follows: RNP complexes were prepared in Lonza P3 Nucleofector Solution by mixing 82 µL of P3 solution, 18 µL of supplement, 3.2 µL Cas9 protein, and 4 µL total sgRNA (2 µL of each sgRNA at 100 µM). Complexes were gently mixed and incubated at RT for 15 min before use. iBMDMs were detached using trypsin, washed once in PBS, and counted. For each nucleofection reaction, 5 x 10^5^ cells were pelleted and resuspended directly in the preassembled RNP complex. Cell–RNP mixtures were transferred to Lonza nucleofection cuvettes and nucleofected using the Lonza 4D-Nucleofector system with program DP-148. Immediately after nucleofection, 400 µL of prewarmed complete medium was added, and cells were transferred to 6-well plates containing fresh medium. Cells were incubated for 48 h prior to downstream analysis.

Genomic DNA was extracted from bulk-edited or clonal cell populations using the PureLink Genomic DNA Mini Kit (Thermo Fisher Scientific) according to the manufacturer’s instructions. DNA was eluted in 40 µL nuclease-free water per well of a 6-well plate. Target loci were PCR-amplified using KOD DNA polymerase with the following primers (5′–3′) *Pycard* (ASC) Forward: TGGCAGGAGGAACAGTTAAGC, Reverse: GCAGCAAGAGTAAAAGGTGACC and *Casp1* Forward: AGGTTGGTTTCTTGAAAGGACT, Reverse: CAGGCAGCAAATTCTTTCACCT. PCR amplicons were column-purified and subjected to Sanger sequencing alongside unedited wild-type controls.

CRISPR KO efficiency was estimated using Inference of CRISPR Edits (ICE; Synthego), comparing edited samples to wild-type reference traces to determine indel frequency and predicted knockout rates. Monoclonal derivation was performed by limiting dilution using conditioned medium. Conditioned medium was prepared by collecting supernatant from subconfluent iBMDM cultures and filtering through a 45 µm syringe filter. Edited cells were passed through a 40 µm cell strainer and diluted to 5 cells mL⁻^1^ in conditioned medium. Aliquots of 100 µL were distributed into 96-well plates, yielding approximately 0.5 cells per well. Clones were expanded and screened by genomic PCR and sequencing.

#### Cell stimulation

For experiments studying NLRP3 inflammasome activation, BMDMs were primed using lipopolysaccharide (LPS; from *Escherichia coli* O26:B6, 1 µg mL^-1^, 4 or 6 h as indicated) in DMEM (10% v/v FBS, 100 U mL^-1^ penicillin, 100 µg mL^-1^ streptomycin). Subsequently, the media was replaced with serum-free DMEM (100 U mL^-1^ penicillin, 100 µg mL^-1^ streptomycin). NLRP3 was then activated by addition of either nigericin (10 µM, 1 h), ATP (5 mM, 1 h), L-Leucyl-L-Leucine methyl ester (LLOME, 1 mM, 1 h), silica (300 µg mL^-1^, 2 or 4 h, as indicated), imiquimod (75 or 150 µM, 2 h) or appropriate vehicle control. NLRP3 stimuli dose-response experiments were performed using concentrations of nigericin (0.1-10 μM), imiquimod (25-200 μM) and silica (1-3000 µg mL^-1^). When included, NLRP3 inhibitors (MCC950 (26), 10 µM) or caspase-1 inhibitors (VX-765, 10 µM) were added 15 min prior to addition of NLRP3 activators. Imiquimod treatment (75 μM, 2 h) was also performed with a monensin (10 μM, 2 h) co-treatment. K^+^ efflux-independent NLRP3 stimuli CL097 (75 μM, 2 h) and forchlorfenuron (75 μM, 2 h) were also used.

For experiments where NLRP3 was activated following non-canonical caspase-11 inflammasome activation, BMDMs were primed using Pam3CSK4 (100 ng mL^-1^, 4 h) in DMEM (10% v/v FBS, 100 U mL^-1^ penicillin, 100 µg mL^-1^ streptomycin). Subsequently, the non-canonical inflammasome was activated via lipofectamine 3000-mediated transfection of LPS (2 µg mL^-1^) or incubated with lipofectamine 3000 alone (0.25 µL/100 µL) or with LPS without transfection reagent (extracellular LPS, 2 µg mL^-1^) for 20 h.

Cell lysates were collected for western blotting and cell supernatants for determination of IL-1β release and pyroptosis. IL-1β release was determined by ELISA (R&D systems, DY401) and pyroptosis was determined by lactate dehydrogenase (LDH) release (Promega, G1780) according to the manufacturer’s instructions.

### Western blotting

Cell lysates were generated by lysis in buffer (50 mM Tris/HCl, 150 mM NaCl, 1% v/v Triton X-100 (Tx100), pH 7.3) containing protease inhibitor cocktail. Where stated, total cell lysates (combined cell lysates and supernatants) were generated by directly adding protease inhibitor cocktail and 1% v/v Tx100 to the well. Lysates were mixed with 5X Laemmli buffer (to a final concentration of 60 mM Tris pH 6.8, 10% v/v glycerol, 5% v/v β-mercaptoethanol, 2% w/v SDS, 0.05% w/v bromophenol blue) and boiled at 95°C for 5 min. For ASC oligomerisation assays, total cell lysates were separated into Tx100-soluble and -insoluble fractions by centrifugation at 6800 *x g* (20 min, 4°C). Tx100-insoluble fractions were crosslinked by incubation with disuccinimidyl suberate (2 mM, 30 min, RT). Crosslinked pellets were further centrifuged at 6800 *x g* (20 min, 4°C) and eluted in Laemmli buffer. Lysates were separated by Tris-glycine SDS PAGE and transferred onto nitrocellulose or PVDF membranes using a semi-dry Trans-Blot Turbo system. Membranes were blocked in either 5% w/v milk or 5% w/v BSA in PBS-T (0.1% v/v Tween-20 in PBS) before overnight incubation with primary antibodies at 4°C. Specific antibodies used for western blotting were: mouse anti-NLPR3 (Adipogen, Cat#AG-20B-0014-C100), mouse anti-RFP (Proteintech, Cat#6g6), rabbit anti-mouse ASC (Cell signalling technology, Cat#67824), rabbit anti-caspase-1 (Abcam, Cat#ab179515), goat anti-mouse IL-1β (R&D systems, Cat#AF-401), rabbit anti-mouse GSDMD (Abcam, Cat#ab209845) and β-actin-HRP (Sigma, Cat#A3854). Membranes were washed three times in PBS-T before incubation with appropriate HRP-conjugated secondary antibodies in 5% w/v BSA in PBS-T at RT for 1 h. After a further three washes in PBS-T, chemiluminescence was visualised using ECL prime (GE healthcare) and a G:box Chemi XX6 (Syngene).

### Immunofluorescence

WT and NLRP3-mScarlet-I BMDMs were seeded out overnight onto 13 mm glass coverslips at a density of 1 x 10^6^ mL^-1^ (500 µL). BMDMs were primed with LPS and stimulated as described above. After stimulation BMDMs were washed once with ice cold PBS (+Ca^2+^/Mg^2+^) and fixed using paraformaldehyde (PFA, 4% w/v in PBS, stabilised with methanol (Sigma, 252549)) at RT for 10-15 min.

For experiments where only NLRP3-mScarlet-I and ASC were labelled, coverslips were washed three times with PBS before permeabilisation and blocking in 1% w/v BSA, 0.1% v/v Triton X-100 in PBS at RT for 1 h. Coverslips were incubated with specific primary antibodies targeting RFP and ASC as described in table 1 in 1% w/v BSA, 0.1% v/v Triton X-100 in PBS at 4°C overnight. Following three washes in PBS, coverslips were incubated with appropriate species-specific highly cross-adsorbed Alexa-Fluor-conjugated secondary antibodies in 1% w/v BSA, 0.1% v/v Triton X-100 in PBS at RT for 1 h. Coverslips were washed three times in PBS before incubation with DAPI (1 µg mL^-1^, 10 min) and washed three times with dH_2_O. Coverslips were air dried and mounted in pro-long gold onto slides before imaging via confocal microscopy.

**Table 1:** Antibodies used for immunofluorescence.

| Target | Host | Secondary used | Clone | Catalogue # | Supplier | Dilution |
| --- | --- | --- | --- | --- | --- | --- |
| RFP (IF) | Rat | Donkey anti-rat alexa-594 (Thermo, A-21209) | 5F8 | 5f8 | Proteintech | 1:250 |
| ASC | Rabbit | Donkey anti-rabbit alexa-488 (Thermo, A-21206) | D2W8U | 67824 | Cell signalling technology | 1:500 |
| ASC-Alexa-fluor488 | Rabbit |  | D2W8U | 17507 | Cell signalling technology | 1:500 |
| PI4P | Mouse IgM | Goat anti-mouse IgM 647 (Thermo, A21238) | PI4-2 | Z-P004 | Echelon biosciences | 1:100 |
| LAMP1 | Rabbit | Donkey anti-rabbit alexa-647 (Thermo, A-31573) | EPR21026 | ab208943 | Abcam | 1:500 |
| TGN38 | Sheep | Donkey anti-sheep alexa-647 (Thermo, A-21448) | Polyclonal | AHP499G | Bio-Rad | 1:100 |
| EEA1 | Mouse | Donkey anti-mouse alexa-647 (Thermo, A32787) | 14/EEA1 | 610457 | BD biosciences | 1:100 |
| γ-tubulin | Mouse | Donkey anti-mouse alexa-647 (Thermo, A32787) | GTU-88 | T6557 | Sigma | 1:500 |

For experiments examining organelle localisation of the NLRP3 inflammasome, different immunostaining protocols were performed as required to preserve required epitopes. For labelling TGN38/ASC/RFP, EEA1/ASC/RFP and γ-tubulin/ASC/RFP, coverslips were processed as described above with antibodies described (Table 1). For labelling PI4P/ASC/RFP, coverslips were washed three times with PBS (+ 50 mM NH_4_Cl) before permeabilisation with 20 µM digitonin in PIPES buffer (20 mM PIPES pH 6.8, 137 mM NaCl, 2.7 mM KCl) at RT for 10 min. Coverslips were then blocked in PBS (+ 50 mM NH_4_Cl) containing 5% v/v goat serum at RT for 1 h before incubation with specific primary antibodies (Table 1) in PBS (+ 50 mM NH_4_Cl) containing 5% v/v goat serum at 4°C overnight. Following three washes in PIPES buffer, coverslips were incubated with appropriate species-specific highly cross-adsorbed Alexa-Fluor-conjugated secondary antibodies in PBS (+ 50 mM NH_4_Cl) containing 5% v/v goat serum at RT for 1 h. Coverslips were washed three times in PIPES buffer before incubation with DAPI (1 µg mL^-1^, 10 min) and washed three times with dH_2_O. Coverslips were then post-fixed by incubation in 2% w/v PFA in PBS at RT for 5 min, washed three times in PBS (+ 50 mM NH_4_Cl), and washed three times with dH_2_O. Coverslips were air dried and mounted in pro-long gold onto slides before imaging via confocal microscopy. For labelling LAMP1/ASC/RFP, coverslips were washed three times with PBS (+ 50 mM NH_4_Cl) before permeabilisation with 20 µM digitonin in PIPES buffer at RT for 10 min. Coverslips were then blocked in PBS (+ 50 mM NH_4_Cl) containing 5% v/v donkey serum at RT for 1 h before incubation with specific primary antibodies (Table 1), excluding ASC, in PBS (+ 50 mM NH_4_Cl) containing 5% v/v donkey serum at 4°C overnight. Following three washes in PIPES buffer, coverslips were incubated with appropriate species-specific highly cross-adsorbed Alexa-Fluor-conjugated secondary antibodies in PBS (+ 50 mM NH_4_Cl) containing 5% v/v donkey serum at RT for 1 h. Coverslips were washed three times in PIPES buffer and then post-fixed by incubation in 2% w/v PFA in PBS at RT for 5 min. Coverslips were washed three times in PBS (+ 50 NH_4_Cl) and then blocked again in PBS (+ 50 mM NH_4_Cl) containing 5% v/v donkey serum and 0.1% v/v Tx100 for 1 h. Coverslips were washed three times with PIPES buffer before incubation with anti-ASC directly conjugated to Alexa-Fluor-488 in PBS (+ 50 mM NH_4_Cl) containing 5% v/v donkey serum and 0.1% v/v Tx100 at 4°C overnight, or RT for 3 h. Coverslips were washed three times in PIPES buffer before incubation with DAPI (1 µg mL^-1^, 10 min) and washed three times with dH_2_O. Coverslips were air dried and mounted in pro-long gold onto slides before imaging via confocal microscopy. Genotype and secondary-only controls were used for all stains to confirm primary antibody specificity.

### Confocal microscopy

Confocal microscopy images were acquired using a 63×/1.40 HCX PL Apo objective on a Leica TCS SP8 AOBS upright confocal microscope with LAS X software (v3.5.2.18963). To prevent interference between channels, lasers were excited sequentially for each channel. A blue diode laser (405 nm) and white light laser (set to excite at 488, 569, 594 and 647 nm) were used, with hybrid and photon-multiplying tube detectors with detection mirror settings set appropriately. Laser powers used for endogenous NLRP3-mScarlet-I were between ∼15-20% and with RFP antibody amplification between ∼8-10%. Z-stacks were acquired with 0.3 µm steps between Z sections. Images were acquired from 2-5 fields of view from each independent experiment.

### Organelle colocalisation analysis

For organelle colocalisation analysis, NLRP3/ASC specks were manually assessed, and speck signal that was determined to be immediately adjacent to, or overlapping with, signal for each organelle label in the XYZ plane was classed as ‘associated’. All specks from each FOV within each biological repeat were combined to determine the percentage of specks associated. The shortest distance between the ASC speck and MTOC (γ-tubulin punctum) signal in the XYZ plane was manually determined using Pythagoras’ theorum and FIJI.

### Single Molecule Localisation Microscopy (dSTORM)

Wild-type and NLRP3-mScarlet-I homozygote BMDMs were plated at 0.5 x 10^6^ cells mL^-1^ (200 µL) into µ-Slide 8 well glass bottom chamber slides (Ibidi: 80827) and left to adhere overnight. BMDMs were primed with LPS (1 µg mL^-1^, 6 h) in DMEM 10% v/v FBS, 100 U mL^-1^ penicillin, 100 µg mL^-1^ streptomycin). After LPS priming, cells were pre-incubated in serum-free DMEM (100 U mL^-1^ penicillin, 100 µg mL^-1^ streptomycin) with the caspase-1 inhibitor VX-765 (10 μM, 15 min), followed by treatment with the NLRP3 activators nigericin (10 μM, 1 h) and imiquimod (150 μM, 1 h).

Cells were washed once with ice cold PBS (+Ca^2+^/Mg^2+^) then fixed with 4% w/v PFA for 10 min at RT before blocking for 30 min in blocking solution (5% w/v BSA and 0.1% v/v Triton X-100 in PBS). Fixed cells were incubated with primary conjugated nanobodies diluted in blocking solution overnight at 4°C. The specific primary nanobodies used were AlexaFluor 647-conjugated FluoTag®-X2 anti-mScarlet-I (1:4000; NanoTag; N1302-AF647-L) and an Atto-488-conjugated anti-ASC (1:1000; produced in house using published sequences (23) and conjugated in house with an Atto-488 NHS ester: Sigma-Aldrich; 41698). Cells were washed in PBS 0.1% v/v Tween-20 followed by post-fixing in 4% w/v PFA before 3x washes in PBS. Samples for two-colour dSTORM images were immersed in 0.22 μm-filtered OxEA imaging buffer (50 mM β-Mercaptoethylamine hydrochloride (MEA; 30078; Sigma Aldrich), 3% v/v OxyFluor™ (OF; Oxyrase Inc.), 20% v/v sodium DL-lactate solution (L4263; Sigma Aldrich) in DPBS, pH adjusted to 8–8.5 with NaOH; (48)). Images were acquired sequentially, firstly by excitation with the 647 nm laser (25% laser power, 7,500 frames; 11 ms exposure time; 120 electron multiplier gain), followed by excitation with the 488 nm laser (100% laser pumping for 2 s followed by acquisition using 55% laser power; 15,000 frames, 11 ms exposure time; 120 electron multiplier gain). For both colours a 405 nm laser (5%) was used after the first 1,000 frames to boost fluorophore recovery.

Single molecule localisation and image reconstruction was performed using ThunderSTORM (49) within the FIJI software. Raw images were filtered using a wavelet filter B-spline method (order 3; scale 2) to remove noise and enhance single fluorophore blinking events. A maximum localisation method (peak intensity threshold = 2 x SD of F1 wavelet; 8-neighbourhood connectivity) was used to determine initial event detection. Subpixel event localisation was calculated using an integrated Gaussian point-spread function and maximum likelihood estimator (fitting radius = 5 pixels; sigma = 1.6 pixels). STORM events were filtered in the 647 channel according to the following criteria: frame >100, intensity >125 photons and sigma between 100 and 180. STORM events in the 488 channel were filtered as follows: frame >500, intensity >300 photons and sigma 30-175. Distinct filtered events detected within 50 nm of each other or within 20 frames of the initial detection were merged to adjust for re-blinking events (as previously published; (50)).

### dSTORM Image Analysis

To analyse NLRP3-mScarlet-I and ASC speck area and roundness, binary masks of dSTORM images were generated using ImageJ’s “Make Binary” function. Circular annotations of 1.75 μm diameter were positioned around ASC-positive specks, within which the area and roundness of NLRP3-mScarlet-I- and ASC-positive signal were measured.

To analyse the distribution of NLRP3-mScarlet-I and ASC signal relative to the centre of the speck, radial averaging using a previously reported ImageJ macro (24) was performed on 2 µm^2^ square annotations manually positioned around the centre of ASC-positive specks. Radial intensity profiles were generated using the “Plot Profile” function in ImageJ. Averaged radial intensity profiles were normalised to the modal value in each condition. To calculate the FWHM of speck radial intensity profiles, nonlinear regression was performed in GraphPad Prism to fit the radial intensity profiles of individual NLRP3-mScarlet-I and ASC specks with a Gaussian model, from which the standard deviation (σ) was extracted. FWHM values were calculated using FWHM = 2√(2 ln 2)·σ.

### In vivo peritoneal inflammation model/Tissue preparation for flow

WT, heterozygous or homozygous NLRP3-mScarlet-I mice were administered with LPS (10 mg kg^-1^, from *Escherichia coli* 0127:B8) intraperitoneally (27). Four hours after injection, mice were stably anesthetised with 2-3% isoflurane in a 70:30% mixture of N_2_/O_2_, and the peritoneal cavity was lavaged with 3 mL of RPMI media and blood was collected via cardiac puncture.

Peritoneal lavage was centrifuged at 1500 *x g* for 5 min at 4°C, the supernatant was stored for cytokine content, and the cell pellet was resuspended in 200 µL ice cold RPMI (3% v/v FBS, 1 mM EDTA, 100 U mL^-1^ penicillin, 100 µg mL^-1^ streptomycin, 2 mM L-glutamine) prior to processing for flow cytometry. Blood was centrifuged at 1500 *x g* for 15 min at 4°C, and the supernatant was subsequently centrifuged at 18000 *x g* for 3 min at 4°C. The initial blood cell pellet underwent two rounds of ACK red blood cell lysis, and the resulting cell pellet was resuspended in 200 µL ice cold RPMI containing 3% v/v FBS, 1 mM EDTA (+P/S + glutamine) prior to staining for flow cytometry. The resulting plasma and peritoneal lavage supernatant were assessed for cytokine content by ELISA. Specific ELISA kits used were IL-1β (R&D, DY401), IL-6 (R&D, DY406) and IL-18 (Thermo, BMS618-3).

Tissues were microdissected and placed into ice-cold HBSS (dorsal portion of the skull, brain hemisphere and spleen). A single tibia was taken from each mouse. Lower mandibles and zygomatic arches were removed from skulls. Skulls were then cut from the foramen magnum, along the squamosal suture to the olfactory bulbs. The ventral portion of the skull was removed to allow extraction of the brain. Dorsal portions of the skull were placed in ice-cold HBSS. Dural meninges were peeled from the inner surface of the skullcap and enzymatically dissociated in digestion buffer containing HBSS (H9269, Sigma) with Liberase TL (1 U mL^-1^; Sigma) and DNase I (80 U mL^-1^; Sigma). Dural samples were incubated at 37°C for 30 min. Samples were resuspended and passed through a 70 μm cell strainer. The digestion process was neutralised by addition of 10x the volume of FACS buffer (0.5% w/v BSA in PBS). Dural suspensions were pelleted (450 *x g*, 5 min) and resuspended in FACS buffer before staining.

Brains were split into hemispheres, with one hemisphere placed into ice-cold HBSS. Brain hemispheres were dounced 20 times in HBSS (H9394, Sigma) to ensure sufficient tissue dissociation. Suspensions were passed through a 70 μm cell strainer and pelleted (300 *x g*, 5 min). Cell pellets were resuspended in 30% Percoll and centrifuged (900 *x g*, 25 min) to remove myelin and debris. The resulting pellet was washed with FACS buffer.

Spleens were cut into small pieces, incubated with digestion buffer at 37°C for 30 min and pushed through a 70 μm cell strainer. The digestion process was neutralised by addition of 10x the volume of FACS buffer. Splenic suspensions were pelleted (450 *x g*, 5 min) and RBCs lysed by incubation in ACK lysis buffer (ThermoFisher). The lysis process was neutralised by addition of 10x the volume of FACS buffer. Splenic suspensions were pelleted (450 *x g*, 5 min), washed and resuspended in FACS buffer.

Bone marrow was collected by cutting the tip of the femur and briefly centrifuging (1000 *x g*, 1 min) the bone marrow into a cell pellet. RBCs were lysed by incubation in ACK lysis buffer before resuspending the pellet in FACS buffer before staining.

### Flow Cytometry

Surface staining was performed by incubating cell suspensions for 60 min at 4°C in FACS buffer (1xPBS/0.5% w/v BSA) with cocktails of antibodies (listed in Table 2), rat anti-mouse CD16/32 (1:100 FcR block) to reduce nonspecific antibody binding and viability dye eFluor™ 780 (1:4000; Invitrogen™ eBioscience™) to exclude dead cells. Samples were washed with FACS buffer, fixed with 4% v/v PFA and resuspended in FACS buffer. Flow cytometry was performed using an Aurora spectral flow cytometer (Cytek Biosciences) and data were analysed with FlowJo (v10; BD Biosciences). Dimensionality reduction was implemented on single livedead^-^ CD45^+^ concatenated and downsampled populations via the tSNE algorithm and clustering was performed with FlowSOM v4.1.0.

**Table 2.** Flow cytometry antibodies.

| Target | Host | Conjugate | Clone | Catalogue # | Supplier | Dilution |
| --- | --- | --- | --- | --- | --- | --- |
| NK1.1 | Mouse | BUV395 | PK136 | 564144 | BD Bioscience | 1:600 |
| CD62L | Rat | BUV496 | MEL-14 | 364-0621-80 | Invitrogen | 1:1000 |
| F4/80 | Rat | BUV563 | T45-2342 | 749284 | BD | 1:400 |
| CD19 | Rat | BUV661 | eBio1D3 | 376-0193-82 | ThermoFisher | 1:500 |
| CD11b | Rat | BUV737 | M1/70 | 612800 | BD | 1:1000 |
| CD45 | Rat | BUV805 | I3/2.3 | 752415 | BD | 1:400 |
| CD64 | Mouse | BV421 | X54-5/7.1 | 139309 | Biolegend | 1:200 |
| CD44 | Rat | BV510 | IM7 | 103044 | Biolegend | 1:500 |
| CD4 | Rat | BV605 | RM4-5 | 100548 | Biolegend | 1:600 |
| CX3CR1 | Mouse | BV650 | SA011F11 | 149033 | Biolegend | 1:200 |
| MHCII (I-A/I-E) | Rat | BV650 | M5/114.15.2 | 107641 | Biolegend | 1:1500 |
| CD11c | A Hamster | BV785 | N418 | 117336 | Biolegend | 1:400 |
| CD3 | Rat | SB550 | 17A2 | 100259 | Biolegend | 1:200 |
| Ly6C | Rat | PerCP Cy5.5 | HK1.4 | 128012 | Biolegend | 1:600 |
| CD163 | Rat | PE-Cy7 | S14049I | 155320 | Biolegend | 1:750 |
| CD8a | Rat | APC | 53-6.7 | 100712 | Biolegend | 1:400 |
| Ly6G | Rat | AF700 | 1A8 | 127621 | Biolegend | 1:300 |
| PD-1 | Rat | APC-Fire810 | 29F.1A12 | 135251 | Biolegend | 1:200 |
| FcR Block | Rat | Purified | 93 | 101302 | Biolegend | 1:100 |

### Cranial window surgery

The cranial window procedure was performed as previously described (51, 52). Briefly, anaesthesia was induced using 5% isoflurane and maintained at 2% isoflurane in a 70:30% mixture of N_2_/O_2_. Constant temperature was maintained at 37 °C using a feedback-controlled heating pad set (Oxford Optronix, United Kingdom). The surgical site on the scalp was shaved and sterilized with Videne (EcoLab, United States), and local anaesthetic was applied (EMLA cream, AstraZeneca, United Kingdom). Buprenorphine (50 μg kg^-1^, Vetergesic, United Kingdom) analgesic was injected subcutaneously at the beginning of the surgery. A midline incision was performed on the scalp to expose the underlying skull. A metal plate was fixed to the skull with cyanoacrylate glue (CP-2 chamber plates, Narishige, Japan). Using a high speed microdrill, a 4 mm craniotomy was performed at a centroid located 4 mm lateral to the midline suture and 3 mm below the coronal suture. Once exposed, the dura was washed with sterile saline. A #0 4 mm glass coverslip (CS-4R, Biochrom) was placed over the craniotomy and glued in place. Dental cement (Super-Bond Universal dental cement, Sun Medical, Japan) was applied to secure the window and head plate. Post-surgery, the mice were placed on a heating pad for recovery.

### Intravital imaging

High-resolution intravital confocal imaging of the exposed pial surface was performed using an upright Leica SP8 Multiphoton microscope (Leica Microsystems, Germany) equipped with a HC FLUOTAR L 25×/0.95 NA water-dipping objective. Images were acquired using Leica LAS X software.

Prior to placement of the cranial window coverslip, mice were administered with intraperitoneal LPS (from *Escherichia coli* 0127:B8). 10 mg kg^-1^ was administered for experiments imaging at 6 h. For 12 and 24 h timepoints, 1 mg kg^-1^ LPS was administered at 0 and 12 h, immediately after imaging. NLRP3-expressing cells were visualised using endogenous fluorescence. When stated, hydrazide 633 (Thermo Fisher Scientific; A30634) was administered intravenously (1 mg kg⁻ ¹) to label the vascular elastin layer (53). CD45-positive leukocytes were labelled using a fluorophore-conjugated anti-CD45 antibody administered intravenously prior to imaging (30-F11 # 35-0451, FITC, Cytek, 5 μg per mouse).

Sequential confocal image acquisition was performed using the 488, 561 and 638 nm laser lines to excite FITC-conjugated anti-CD45 antibody, endogenous mScarlet-I fluorescence and Hydrazide 633, respectively. Fluorescence emission was collected using HyD detectors with sequential scanning enabled to minimise spectral overlap between fluorophores. Images were acquired at 1024 × 1024 pixel resolution (16-bit depth), using a scan speed of 400 Hz, frame averaging of 2. XY images and Z-stacks (1 μm z-step) were collected using identical acquisition settings across all animals within each experimental cohort. Laser power (∼8%), detector gain and pinhole size were maintained constant across all animals within each experiment. To differentiate between autofluorescence and specific NLRP3-mScarlet-I signal, images were also acquired in channels containing no fluorophore. Image processing was performed using Fiji. To reduce noise, images were processed with a median filter (radius of 0.6 pixels) and outliers removed (threshold 60, 1-1.5 pixels) was applied.

### Quantification and statistical analysis

Data are mean ± SEM or median ± IQR as stated in figure legends. Individual data points are shown where possible. Data were assessed for normal distribution using Shapiro-Wilk test. Parametric data were analysed using unpaired t-test, one-way ANOVA followed by Tukey’s, Sidak’s or Dunnett’s post-hoc tests, or two-way ANOVA with uncorrected Fisher’s LSD post-hoc analysis. Non-parametric data were analysed using Mann-Whitney test or Kruskal-Wallis followed by Dunn’s post-hoc test. Concentration-response curves were fitted using a four-parameter logistical model to determine EC_50_ values. Correlations were determined using a simple linear regression. GraphPad Prism (v10) was used for all statistical analysis. Statistical significance was accepted at *p<0.05.

## Data availability

All data are available upon request. dSTORM data is available at nano-org (54).

## Acknowledgements

We thank the Flow Cytometry and Bioimaging core facilities at the University of Manchester. We thank the Biomolecular Analysis core facility for generation of the ASC nanobody.

