## Supplementary Figures for "*In vivo* cellular localisation and nanoscale organisation of NLRP3 inflammasomes"

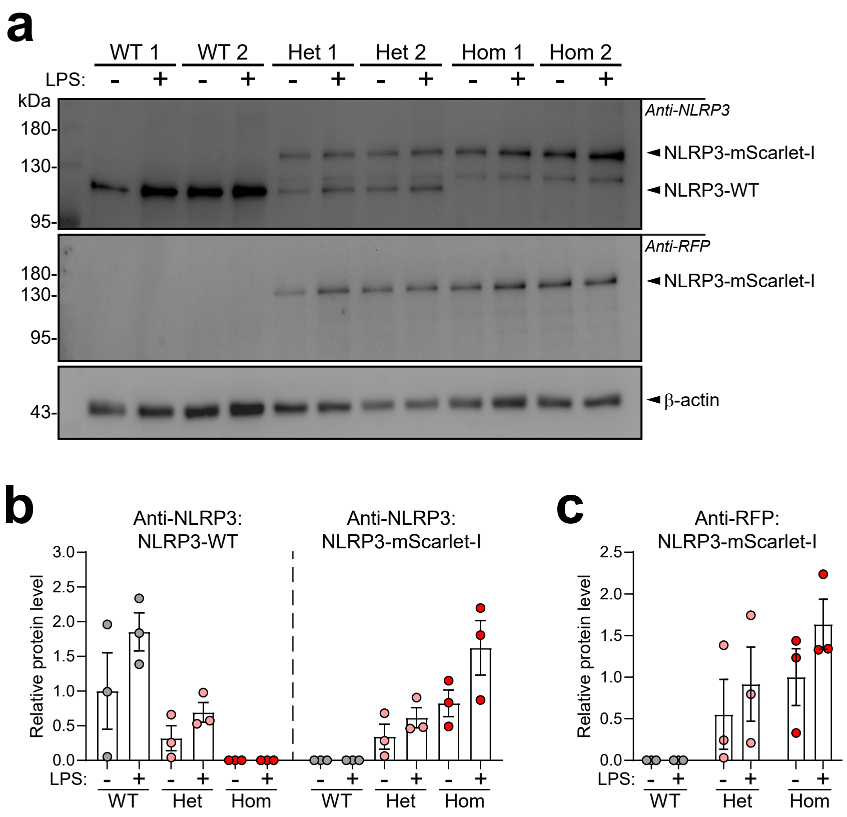


**Supplementary Figure 1. Endogenous NLRP3-mScarlet-I expression in primary mouse peritoneal lavage cells. (a-c)** Peritoneal lavage cells were untreated or primed with LPS (1 µg $mL^{-1}$, 4 h) and (a) cell lysates were blotted for NLRP3 and mScarlet-I (representative from n=3). Densitometry of (b) anti-NLRP3 blots (NLRP3-WT (~115 kDa) and NLRP3-mScarlet-I (~145 kDa)) and (c) anti-RFP blots (NLRP3-mScarlet-I (~145 kDa)), normalised to β-actin (n=3). Data are mean ± SEM. Data were analysed using unpaired t-test (b, not significant, WT LPS vs Hom LPS).

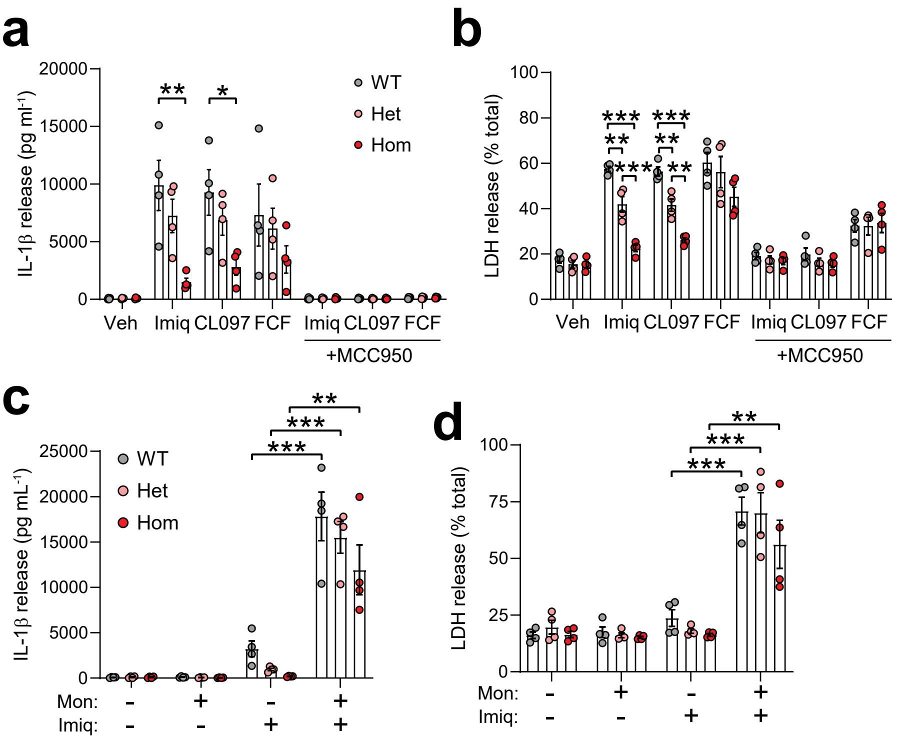


**Supplementary Figure 2. NLRP3-mScarlet-I exhibits reduced sensitivity to K+ efflux-independent stimuli. (a, b)** BMDMs were primed with LPS (1 µg $mL^{-1}$, 4 h) and then treated ± MCC950 (10 µM, 15 min), before addition of imiquimod (Imiq, 75 µM, 2 h), CL097 (75 µM, 2 h), or forchlorfenuron (FCF, 75 µM, 2 h) (n=4). (a) IL-1β and (b) LDH release into the supernatant were measured. **(c, d)** BMDMs were primed with LPS (1 µg $mL^{-1}$, 4 h) and then co-treated with vehicle or imiquimod (75 µM, 2 h) and vehicle or monensin (10 µM, 2 h) (n=4). (c) IL-1β and (d) LDH release into the supernatant were measured. Data are mean ± SEM. Data were analysed using one-way ANOVA followed by Tukey's (a,b) or Sidak's (c,d) post-hoc analysis. *p<0.05, **p<0.01, *** p<0.001.

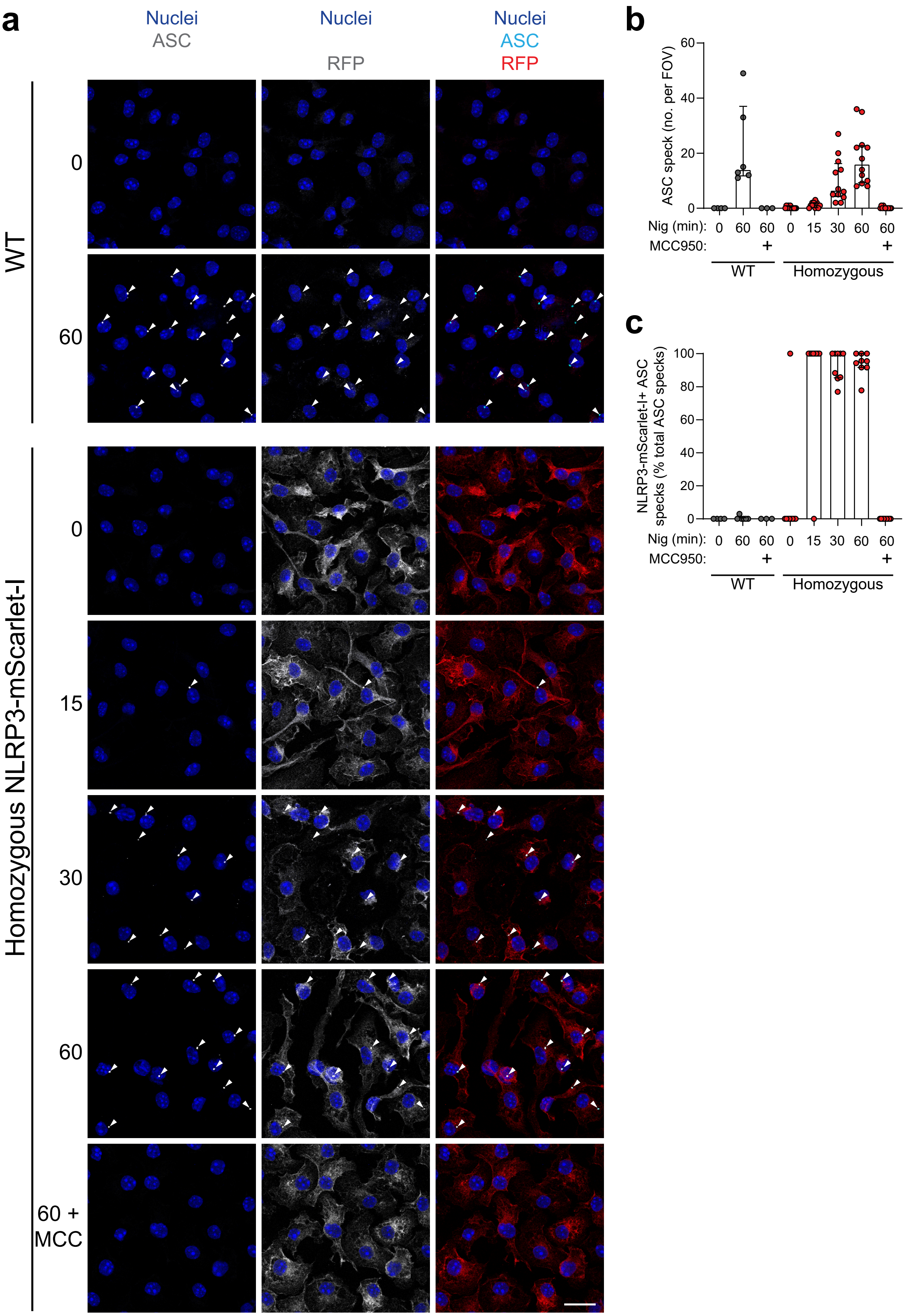
a
Nuclei
ASC
Nuclei
RFP
Nuclei
ASC
RFP
WT
0
60
Homozygous NLRP3-mScarlet-I
0
15
30
60
60 +
MCC
b
ASC speck (no. per FOV)
60
40
20
0
Nig (min): 0 60 60 0 15 30 60 60
MCC950: + +
WT
Homozygous
c
NLRP3-mScarlet-I+ ASC specks (% total ASC specks)
100
80
60
40
20
0
Nig (min): 0 60 60 0 15 30 60 60
MCC950: + +
WT
Homozygous

**Supplementary Figure 3. Time-dependent formation of NLRP3-mScarlet-I specks that colocalised with ASC specks. (a-c)** WT or NLRP3-mScarlet-I BMDMs were primed with LPS (1 μg $ml^{-1}$, 4 h), and then treated ± MCC950 (10 μM, 15 min), before addition of nigericin (10 μM, 0-60 min) (n=2 WT and n=4 Hom – 2-3 FOV from 2-4 biological repeats). (a) Cells were assessed by immunofluorescence for ASC (green) and NLRP3-mScarlet-I using anti-RFP labelling of mScarlet-I (red). A single z-plane is shown. Scale bar = 20 μm. (b) ASC speck number per FOV and (c) % of NLRP3-mScarlet-$I^+$ ASC specks were quantified. Data are median ± IQR.

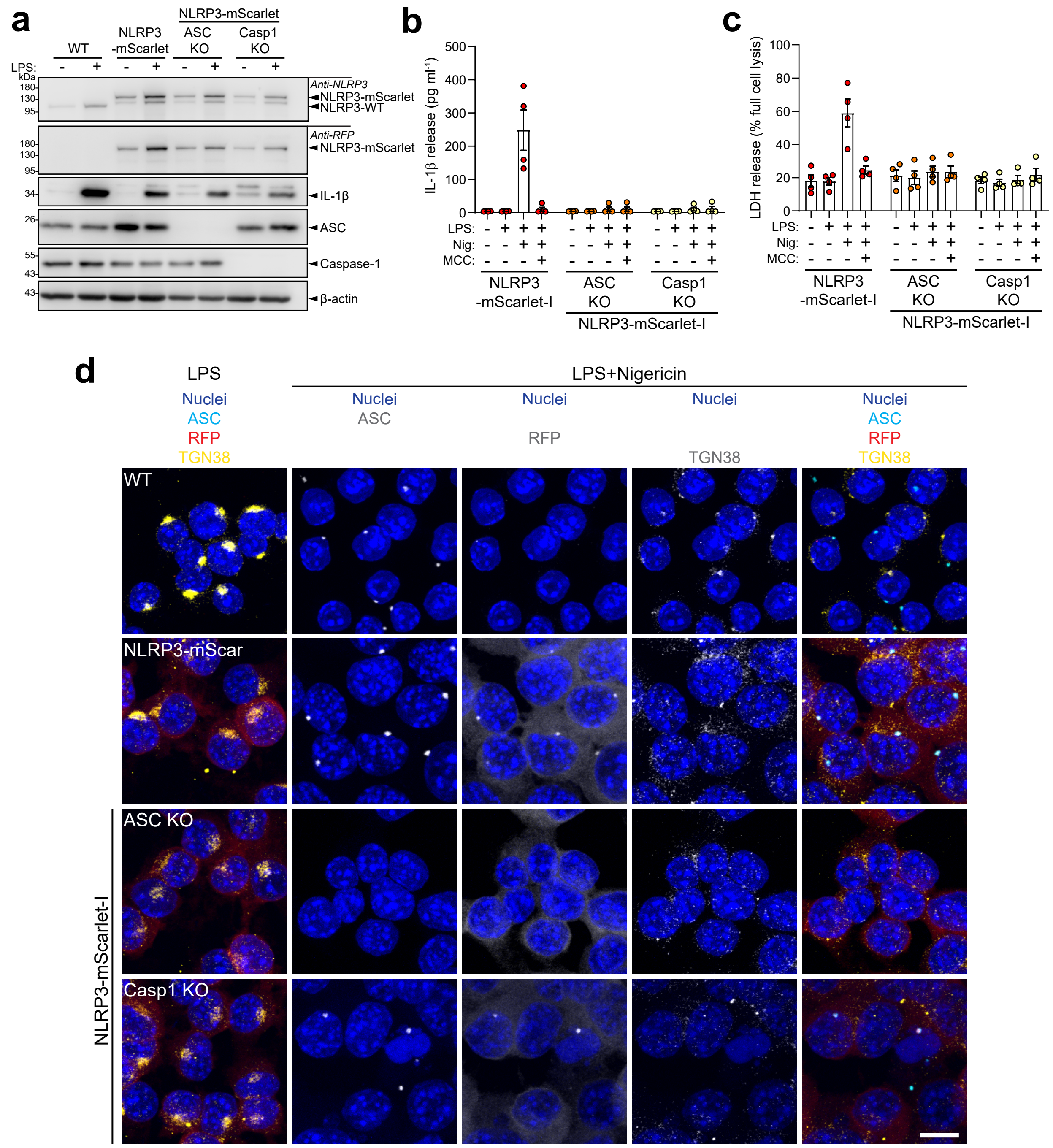


**Supplementary Figure 4. Characterisation of ASC and Casp1 knockout NLRP3-mScarlet-I iBMDMs. (a)** WT, NLRP3-mScarlet-I, ASC knockout (KO) NLRP3-mScarlet-I, or caspase-1 (Casp1) KO NLRP3-mScarlet-I immortalised BMDMs (iBMDMs) were treated ± LPS (1 µg mL$^{-1}$, 4 h). Lysates were blotted for NLRP3, RFP (mScarlet-I), IL-1β, ASC and caspase-1 (representative from n=4). **(b, c)** NLRP3-mScarlet-I, ASC KO NLRP3-mScarlet-I, or Casp1 KO NLRP3-mScarlet-I iBMDMs were treated with LPS (1 µg mL$^{-1}$, 4 h) and then treated ± MCC950 (10 µM, 15 min), before addition of nigericin (10 µM, 1 h) (n=4). (b) IL-1β and (c) LDH release into the supernatant were measured. **(d)** WT, NLRP3-mScarlet-I, ASC KO NLRP3-mScarlet-I, or Casp1 KO NLRP3-mScarlet-I iBMDMs were treated with LPS (1 µg mL$^{-1}$, 4 h) and then treated with VX765 (10 µM, 15 min) before the addition of nigericin (10 µM, 1 h) (n=2). Cells were assessed by immunofluorescence for ASC (cyan), NLRP3-mScarlet-I using anti-RFP labelling of mScarlet (red) and TGN38 (yellow). Maximum intensity projections are shown. Scale bar = 10 µm. Data are mean ± SEM.

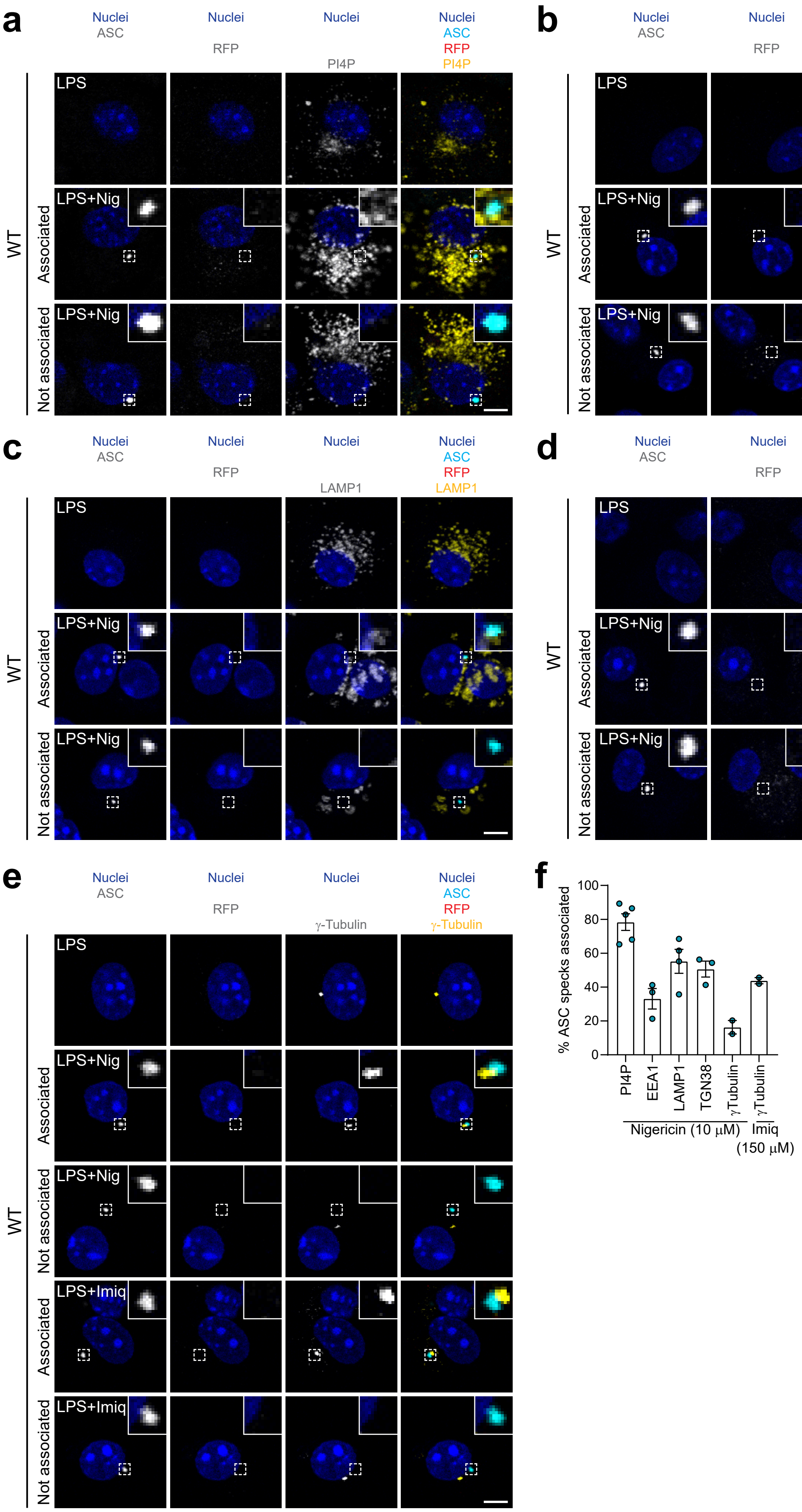

a
Nuclei ASC
Nuclei RFP
Nuclei PI4P
Nuclei ASC RFP PI4P
WT
LPS
LPS+Nig
Associated
Not associated
b
Nuclei ASC
Nuclei RFP
Nuclei EEA1
Nuclei ASC RFP EEA1
c
Nuclei ASC
Nuclei RFP
Nuclei LAMP1
Nuclei ASC RFP LAMP1
d
Nuclei ASC
Nuclei RFP
Nuclei TGN38
Nuclei ASC RFP TGN38
e
Nuclei ASC
Nuclei RFP
Nuclei γ-Tubulin
Nuclei ASC RFP γ-Tubulin
LPS+Imiq
f
% ASC specks associated
0
20
40
60
80
100
PI4P
EEA1
LAMP1
TGN38
γTubulin
γTubulin
Nigericin (10 μM)
Imiq (150 μM)

**Supplementary Figure 5. Subcellular localisation of NLRP3 inflammasomes in WT BMDMs.** Primary WT BMDMs were primed with LPS (1 μg $mL^{-1}$, 6 h), and then treated with nigericin (10 μM, 30 min) or imiquimod (150 μM, 30 min). **(a-e)** Cells were assessed by immunofluorescence for ASC (cyan) and NLRP3-mScarlet-I using anti-RFP labelling of mScarlet (red), as well as (a) PI4P, (b) EEA1, (c) LAMP1, (d) TGN38, (e) γ-tubulin (yellow) (n=2-5 – 3-4 FOV averaged from 2-5 biological repeats). Example NLRP3/ASC specks that were classified as 'associated' or 'not associated' with the respective organelle marker are shown. Single z-planes are shown. Scale bar = 5 μm. (f) Association of ASC specks with each organelle marker. Data are mean ± SEM.

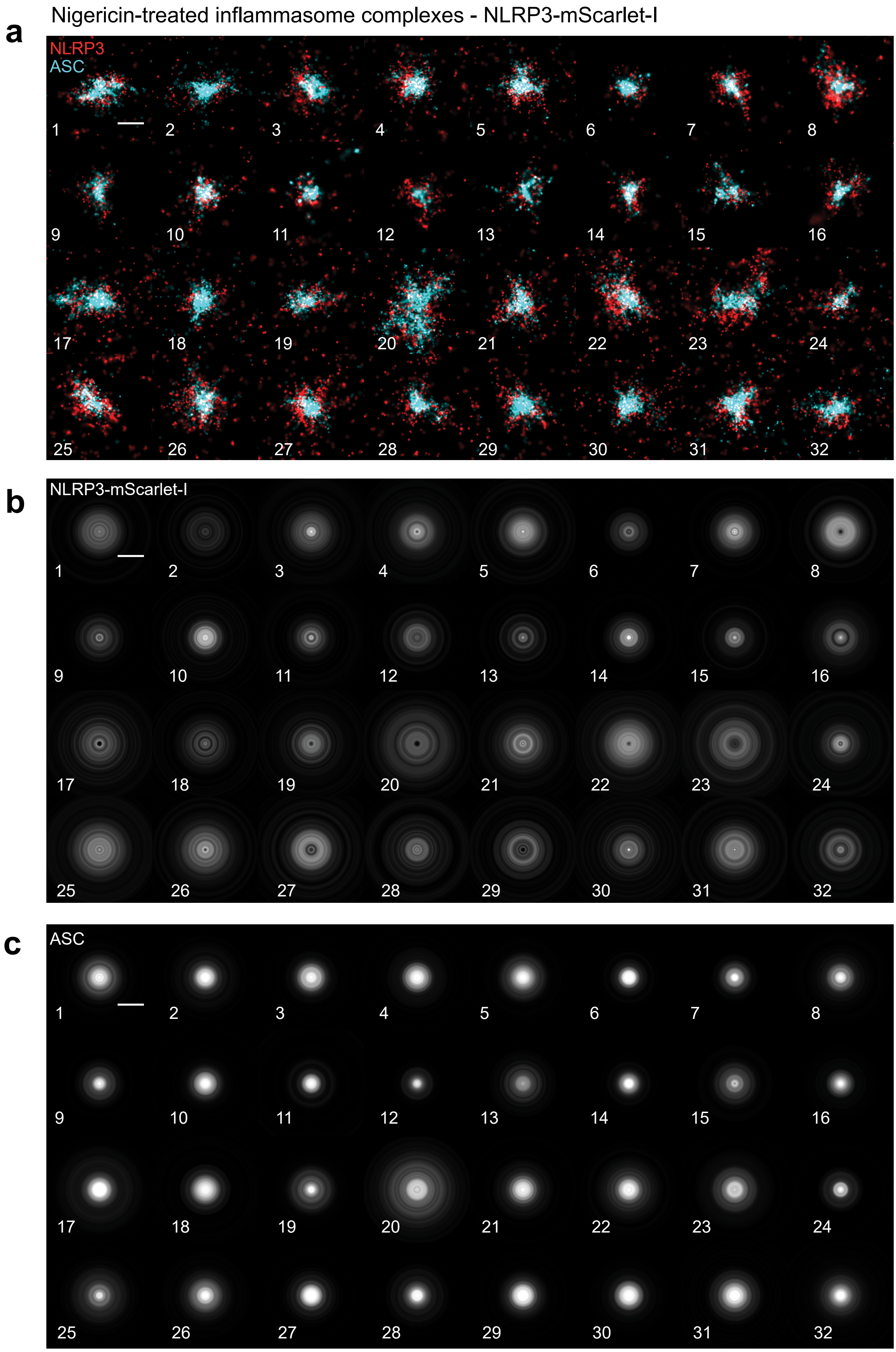
a
Nigericin-treated inflammasome complexes - NLRP3-mScarlet-I
NLRP3
ASC
1 2 3 4 5 6 7 8
9 10 11 12 13 14 15 16
17 18 19 20 21 22 23 24
25 26 27 28 29 30 31 32
b
NLRP3-mScarlet-I
1 2 3 4 5 6 7 8
9 10 11 12 13 14 15 16
17 18 19 20 21 22 23 24
25 26 27 28 29 30 31 32
c
ASC
1 2 3 4 5 6 7 8
9 10 11 12 13 14 15 16
17 18 19 20 21 22 23 24
25 26 27 28 29 30 31 32

**Supplementary Figure 6. Individual dSTORM images and radial averaging images for each nigericin-induced NLRP3 inflammasome complex analysed in NLRP3-mScarlet-I BMDMs.** LPS-primed (1 µg $mL^{-1}$, 4 h) NLRP3-mScarlet-I BMDMs were treated with nigericin (10 µM, 1 h). Each numbered panel represents an individual inflammasome complex analysed. **(a)** dSTORM images of NLRP3 (red) and ASC (cyan). Scale bar = 0.5 µm. **(b,c)** Radial averaging analysis of NLRP3 (b) and ASC (c) from corresponding inflammasome complexes numbered in (a).

Imiquimod-treated inflammasome complexes - NLRP3 mScarlet-I

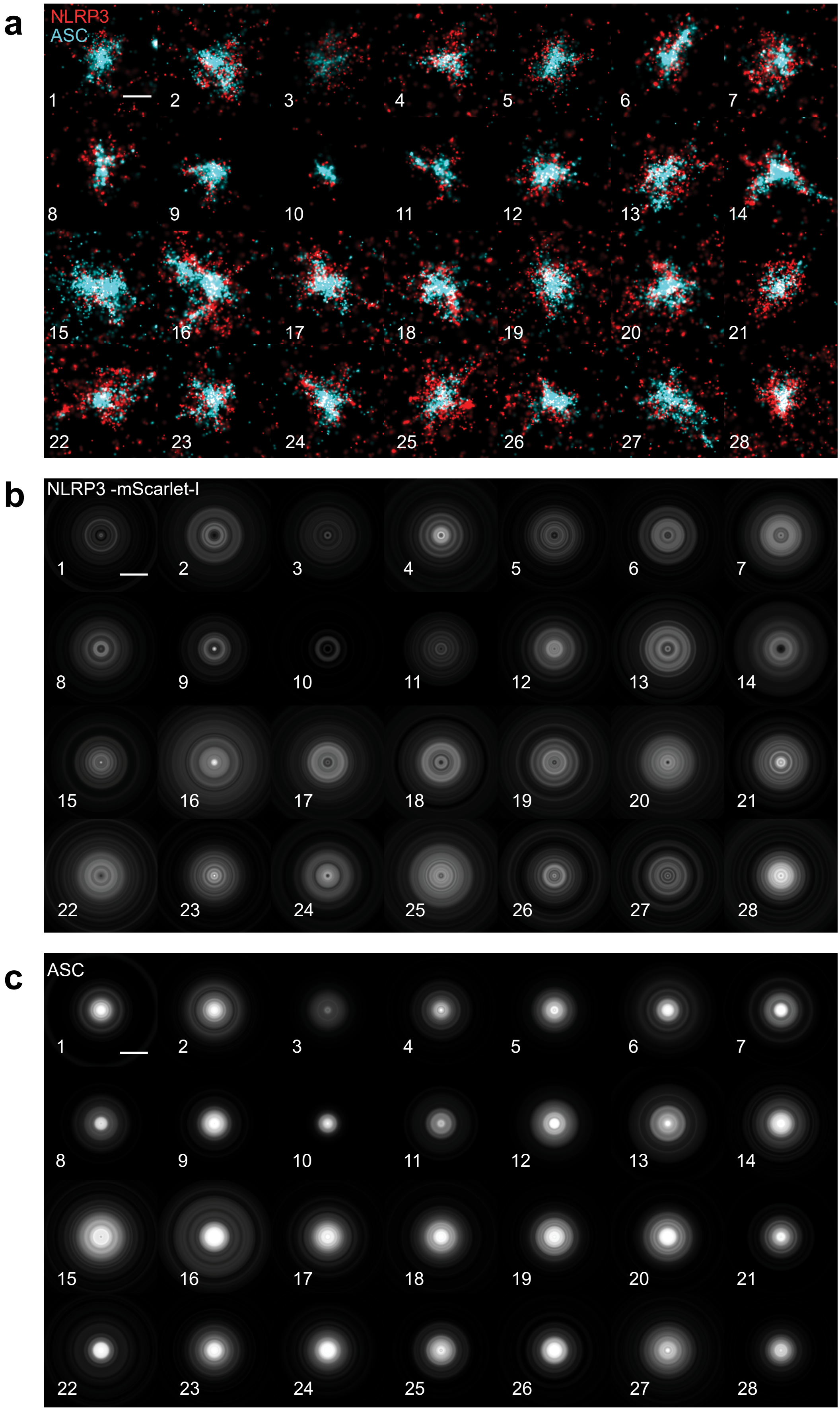

**Supplementary Figure 7. Individual dSTORM images and radial averaging images for each imiquimod-induced NLRP3 inflammasome complex analysed in NLRP3-mScarlet-I BMDMs.** LPS-primed (1 µg $mL^{-1}$, 4 h) NLRP3-mScarlet-I BMDMs were treated with imiquimod (150 µM, 1 h). Each numbered panel represents an individual inflammasome complex analysed. **(a)** dSTORM images of NLRP3 (red) and ASC (cyan). Scale bar = 0.5 µm. **(b, c)** Radial averaging analysis of NLRP3 (b) and ASC (c) from corresponding inflammasome complexes numbered in (a).

Nigericin-treated inflammasome complexes - Wild-type

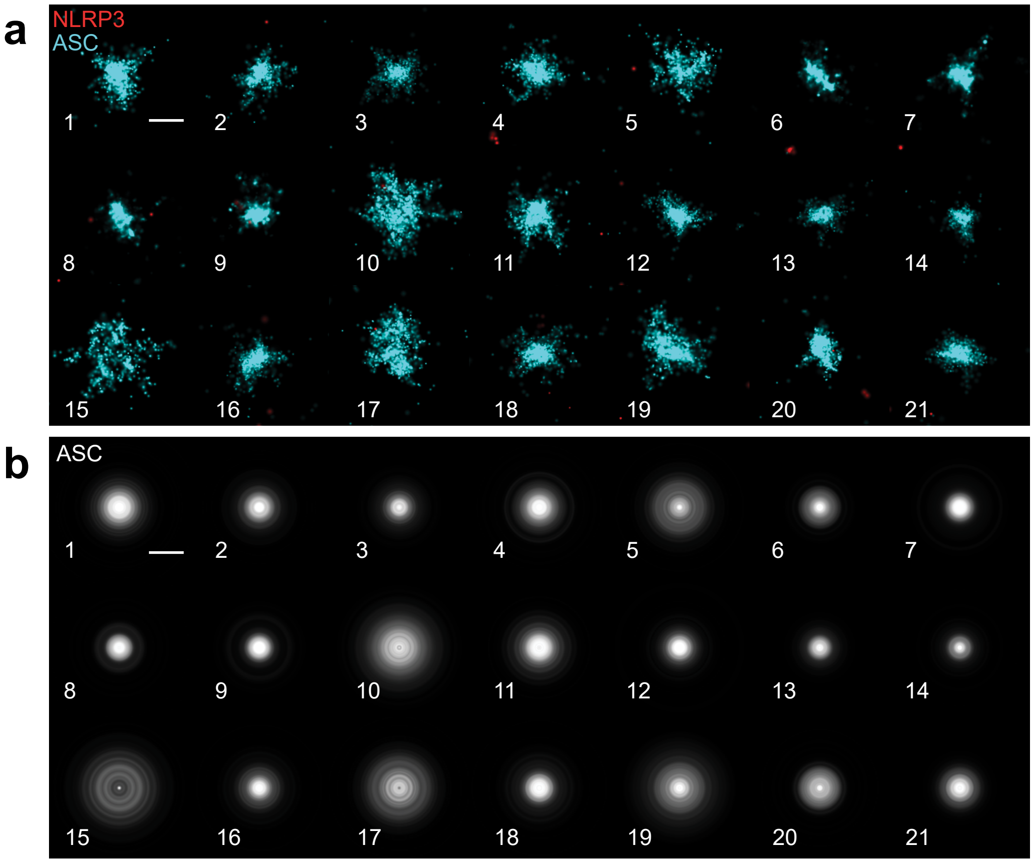


Imiquimod-treated inflammasome complexes - Wild-type

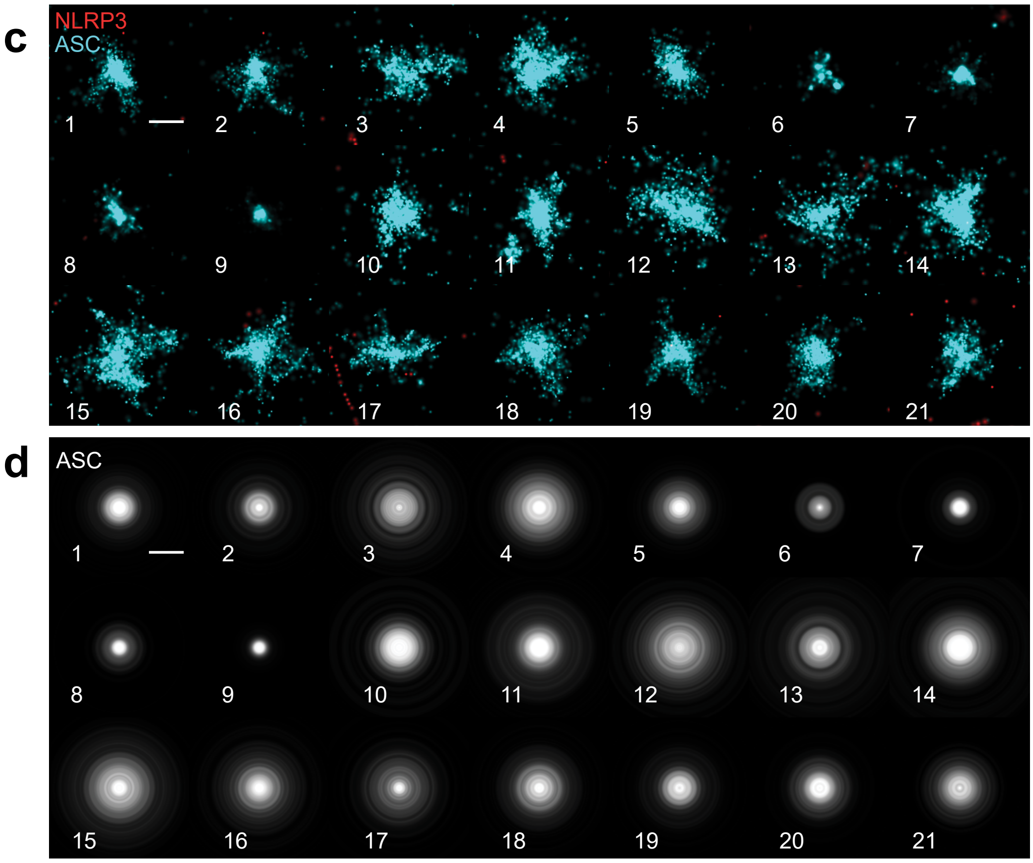

**Supplementary Figure 8. Individual dSTORM images and radial averaging images for ASC specks in WT BMDMs.** LPS-primed (1 µg $mL^{-1}$, 4 h) WT BMDMs were treated with nigericin (10 µM, 1 h) (a-b) or imiquimod (150 µM, 1 h) (c-d). Each numbered panel represents an individual inflammasome complex analysed. **(a)** dSTORM images of ASC (cyan) and mScarlet-signal (red) following nigericin treatment. Scale bar = 0.5 µm. **(b)** Radial averaging analysis of ASC from corresponding inflammasome complexes numbered in (a). **(c)** dSTORM images of ASC (cyan) following imiquimod treatment. Scale bar = 0.5 µm. **(d)** Radial averaging analysis of ASC from corresponding inflammasome complexes numbered in (c).

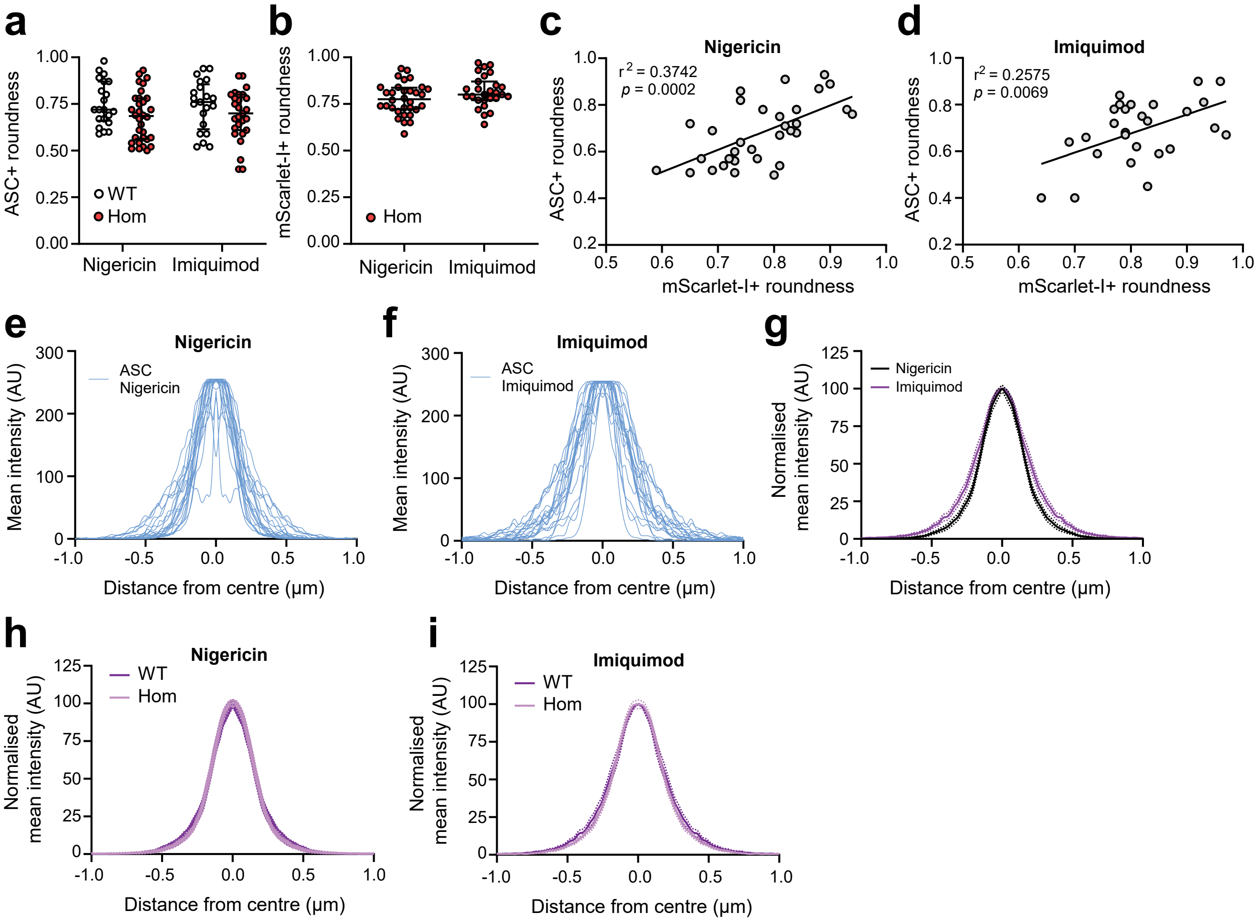


**Supplementary Figure 9. Further analysis of dSTORM images of NLRP3-mScarlet-I and ASC specks.** BMDMs were treated with the same experimental conditions as described in Figure 5. **(a,b)** Roundness of ASC-positive (a) and NLRP3-mScarlet-I-positive (b) staining in inflammasome specks. **(c,d)** Correlations of ASC- and NLRP3-mScarlet-positive roundness in (c) nigericin- and (d) imiquimod-induced specks. **(e-g)** Individual (e) nigericin, (f) imiquimod and (g) mode-normalised average radial intensity profiles of ASC within specks of WT BMDMs. **(h,i)** Mode-normalised average radial intensity graphs comparing ASC distribution in WT and NLRP3-mScarlet-I BMDMs in (h) nigericin- and (i) imiquimod-induced specks. Data in (h) and (i) is replotted from (g) and Fig 5I. Data are median ± IQR. Data were analysed using two-way ANOVA with uncorrected Fisher's LSD post-hoc analysis (a, not significant) or unpaired t-test (b, not significant). Correlations were determined using a simple linear regression.

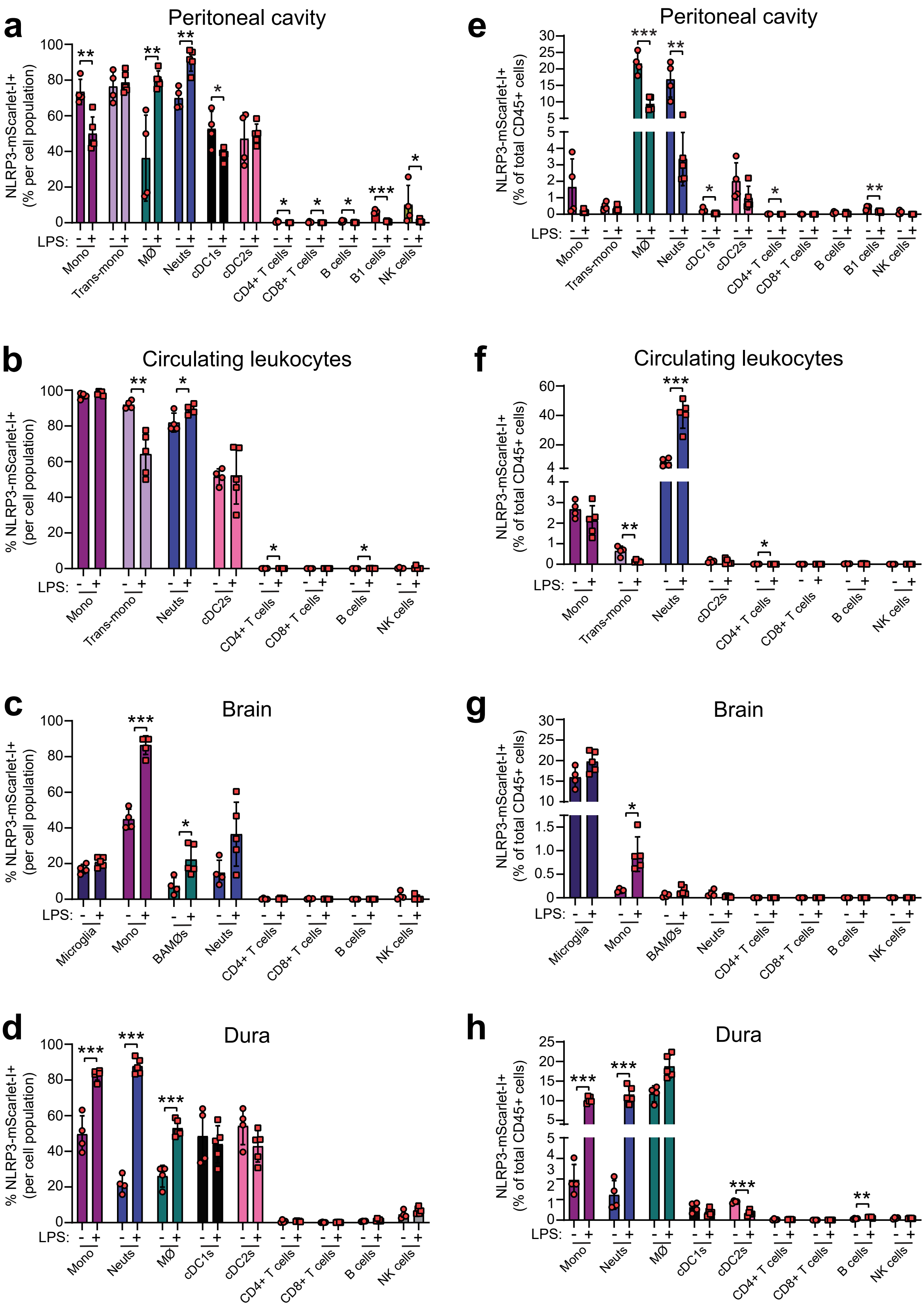
a
Peritoneal cavity
NLRP3-mScarlet-I+ (% per cell population)
LPS:
Mono
Trans-mono
MØ
Neuts
cDC1s
cDC2s
CD4+ T cells
CD8+ T cells
B cells
B1 cells
NK cells
b
Circulating leukocytes
% NLRP3-mScarlet-I+ (per cell population)
c
Brain
Microglia
BAMØs
d
Dura
e
Peritoneal cavity
NLRP3-mScarlet-I+ (% of total CD45+ cells)
f
Circulating leukocytes
g
Brain
h
Dura

**Supplementary Figure 10. Flow cytometry analysis of NLRP3-mScarlet-I expression.** Related to data in Figure 7. **(a-d)** Cells positive for NLRP3-mScarlet-I expression, expressed as a % of each population, in the peritoneum (a), blood (b), brain (c) and dura (d) (n=4-5). (**e-h)** Cells positive for NLRP3-mScarlet-I expression, expressed as a % of total CD45+ cells, in the peritoneum (e), blood (f), brain (g) and dura (h). Data are mean ± SEM. Data were analysed using unpaired t-test (for parametric) or Mann-Whitney test (for non-parametric) (PBS vs LPS per cell type). *$p<0.05$, **$p<0.01$, *** $p<0.001$.

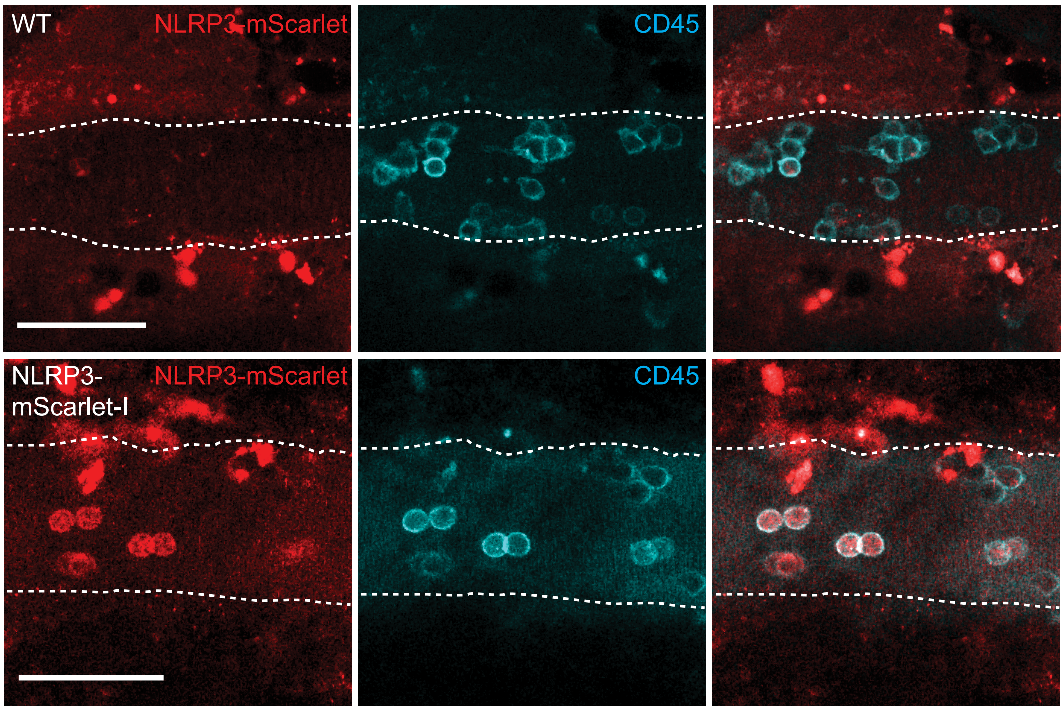


**Supplementary Figure 11. NLRP3-mScarlet-I CD45-positive leukocytes roll on blood vessels following LPS administration.** Intravital imaging of the brain in WT and NLRP3-mScarlet-I littermates following intraperitoneal LPS (10 mg kg$^{-1}$, 6 h). Representative confocal images of NLRP3-mScarlet-I (red) and anti-CD45-FITC (cyan, administrated intravenously). The vasculature is shown by the dotted white line. Scale bar = 50 µm (n=2 WT, n=1 NLRP3-mScarlet-I).

tcttctcaaagattagacaactgcagcctcacctcacacagctgctggaatctctccacaattctgacccacaaccacagccttcg
gaagctgaacctgggcaacaatgatcttggcgatctgtgcgtggtgaccctctgtgaggtgctgaaacagcagggctgcctcctgca
gagcctacagtgagtgtggtttgcctagagcttctcatggggtaggcgagcggggtgctgaggggagggtgaccacgggacaaaagt
cagagtttctctggattaatttgcagttttctgaagagtccaactcaaagcttcttttctgtgttcacaggttgggtgaaatgtacttaaatc
gtgaaacaaaacgtgccttagaagcgctccaggaagaaaagcctgagctgactatagtcttcgagatttcctggGGTGGCTCT
GGCGGAAGCGGCTCCGCTGGCTCTGCTGCCGGATCTGGCGAGTTTGTGAGCAAGGGCGAGGCAG
TGATCAAGGAGTTCATGCGGTTCAAGGTGCACATGGAGGGCTCCATGAACGGCCACGAGTTCGAGAT
CGAGGGCGAGGGCGAGGGCCGCCCCTACGAGGGCACCCAGACCGCCAAGCTGAAGGTGACCA
AGGGTGGCCCCCTGCCCTTCTCCTGGGACATCCTGTCCCCTCAGTTCATGTACGGCTCCAGGGCCT
TCATCAAGCACCCCGCCGACATCCCCGACTACTATAAGCAGTCCTTCCCCGAGGGCTTCAAGTGGG
AGCGCGTGATGAACTTCGAGGACGGCGGCGCCGTGACCGTGACCCAGGACACCTCCCTGGAGGA
CGGCACCCTGATCTACAAGGTGAAGCTCCGCGGCACCAACTTCCCTCCTGACGGCCCCGTAATGC
AGAAGAAGACAATGGGCTGGGAAGCGTCCACCGAGCGGTTGTACCCCGAGGACGGCGTGCTGAA
GGGCGACATTAAGATGGCCCTGCGCCTGAAGGACGGCGGCCGCTACCTGGCGGACTTCAAGACC
ACCTACAAGGCCAAGAAGCCCGTGCAGATGCCCGGCGCCTACAACGTCGACCGCAAGTTGGACAT
CACCTCCCACAACGAGGACTACACCGTGGTGGAACAGTACGAACGCTCCGAGGGCCGCCACTCC
ACCGGCGGCATGGACGAGCTGTACAAGtaggcgtggaagcaggaccaccaggtgcctcggtcctgccccaagtcct
gccccaagccccagtgcgcactgctcttcactgctatcaagccctccttcaccatcaggatcacagccgaggctcttctggtatagg
gtctggagcaaaggcttgtgtgggaccaaatatttttcctcacatcgataacgtgaaactgccagaggctgcccttcccatcatatcc
tcagtgggcaaggtgttccctcttggtgacttcatggaaacagcttcaagaaaacgcct

Lower case text – homology arms to Nlrp3 sequences for C terminal targeting

YELLOWCYAN – GGSWaldo flexible linker

RED – mScarlet-I

**Supplementary Figure 12. The allele sequence for linking mScarlet-I to the c-terminus of NLRP3.**

**Supplementary Video 1. Intravital imaging of NLRP3-mScarlet-I-positive cells rolling within the vasculature 6 h after 10 mg/kg LPS.** Related to Fig 8a-c and Supplementary Fig 11. WT and NLRP3-mScarlet-I mice had cranial windows inserted and the brain was imaged following intraperitoneal LPS (10 mg $kg^{-1}$, 6 h). **(a)** Representative confocal timelapse of NLRP3-mScarlet-I (561 nm, red) and auto-fluorescent background (488 nm, cyan), with resulting non-specific autofluorescence labelled as white. The vasculature is shown by the dotted white line. Scale bar = 50 μm (n=4). **(b)** Representative confocal timelapse of NLRP3-mScarlet-I (561 nm, red) and auto-fluorescent background (488 nm, cyan) showing NLRP3-mScarlet-I-positive cells rolling on endothelium. Scale bar = 25 μm (n=4). **(c)** Representative confocal timelapse of NLRP3-mScarlet-I (561 nm, red) and anti-CD45 (488 nm, cyan, administered intravenously) showing NLRP3-mScarlet-I-positive cells are CD45-positive. Arrows indicate NLRP3-positive, CD45-positive cells. Scale bar = 25 μm, n=2 WT, n=1 NLRP3-mScarlet-I). Time shown in top left in seconds.

**Supplementary Video 2. Intravital imaging of NLRP3-mScarlet-I-positive cells transitioning from vascular adhesion at 12 h to extravascular accumulation at 24 h after 1 mg/kg LPS.** Related to Fig 8d-g. Intravital imaging of the brain in WT and NLRP3-mScarlet-I littermates following insertion of cranial windows and i.p. injection of LPS (1 mg $kg^{-1}$, administered twice, 12 h apart), imaged at 12 h and 24 h. **(a)** Representative confocal timelapse at 12 h post LPS of NLRP3-mScarlet-I (561 nm, red) and auto-fluorescent background (488 nm, cyan), with resulting non-specific autofluorescence labelled as white. The vasculature is shown by the dotted white line. Scale bar = 25 μm (n=3). **(b)** Representative confocal timelapse at 24 h post LPS of NLRP3-mScarlet-I (561 nm, red) and indicated stains (autofluoresence or anti-CD45; cyan, hydrazide; yellow). Scale bar = 25 μm (n=3). Arrows highlight examples of NLRP3-mScarlet-I positive cells. Time shown in top left in seconds.
